# Feedforward and feedback interactions between visual cortical areas use different population activity patterns

**DOI:** 10.1101/2021.02.08.430346

**Authors:** João D. Semedo, Anna I. Jasper, Amin Zandvakili, Amir Aschner, Christian K. Machens, Adam Kohn, Byron M. Yu

## Abstract

Brain function relies on the coordination of activity across multiple, recurrently connected, brain areas. For instance, sensory information encoded in early sensory areas is relayed to, and further processed by, higher cortical areas and then fed back. However, the way in which feedforward and feedback signaling interact with one another is incompletely understood. Here we investigate this question by leveraging simultaneous neuronal population recordings in early and midlevel visual areas (V1-V2 and V1-V4). Using a dimensionality reduction approach, we find that population interactions are feedforward-dominated shortly after stimulus onset and feedback-dominated during spontaneous activity. The population activity patterns most correlated across areas were distinct during feedforward- and feedback-dominated periods. These results suggest that feedforward and feedback signaling rely on separate “channels”, such that feedback signaling does not directly affect activity that is fed forward.

---

Most brain functions rely on the coordination of activity across multiple areas^1,2^. Activity does not follow a purely feedforward path between brain areas: areas are often reciprocally connected, and signals passed from one area to the next are often processed and fed back^3–6^. Understanding when feedforward and feedback signaling between areas is most dominant, and how these forms of signaling interact, is crucial for improving our understanding of computation in the brain.

Previous studies have attempted to infer feedforward or feedback interactions between areas. One approach for identifying feedforward signaling is to present a stimulus and then compare the timing of neuronal response onsets across areas^7–10^. Similarly, feedback signaling can be inferred by studying time differences in the emergence of some forms of selectivity across areas^11–15^. Other studies have studied feedforward or feedback signaling by measuring activity simultaneously in two areas, and comparing temporal delays in pairwise spiking correlations^16–21^or phase delays in local field potentials (LFP)^22–25^. Most of these studies focused on the activity of pairs of neurons across areas, or aggregate measures of neural activity such as local field potentials.

To understand inter-areal interactions more deeply, it is now possible to record activity from large neuronal populations simultaneously in different cortical areas, and characterize what patterns of population activity are most related across those areas^19,26–33^. This approach has led to new proposals about how activity can be flexibly routed across brain areas (see ref. 34 for a review). In particular, simultaneous multi-area recordings have revealed properties of population activity patterns that are most related across areas in the context of sensory processing^29^, attention^30^, learning^31^, and motor control^32,33^. However, it is unknown how these population activity patterns relate to feedforward or feedback signaling between areas.

Here, we leverage simultaneous recordings of neuronal populations in early and midlevel visual areas (V1-V2 and V1-V4) to examine the temporal dynamics of inter-areal interactions, as well as the population activity patterns involved in those interactions (Fig. 1a). We correlated the population activity across areas at different time delays to infer feedforward and feedback signaling. Interactions were feedforward-dominated (V1 leading V2, and V1 leading V4) shortly after stimulus onset and gradually became feedback-dominated with persistent stimulus drive, as well as during spontaneous activity. Importantly, the population activity patterns involved in feedforward signaling were distinct from those involved in feedback signaling. This indicates that activity patterns in V1 that most affect downstream activity during feedforward processing are not the ones most affected by feedback signaling, suggesting both forms of signaling can co-exist without interference. Our results reveal both the dominant direction of signal flow between areas on a moment-by-moment basis and the population activity patterns involved in feedforward and feedback interactions.

**Figure 1.**
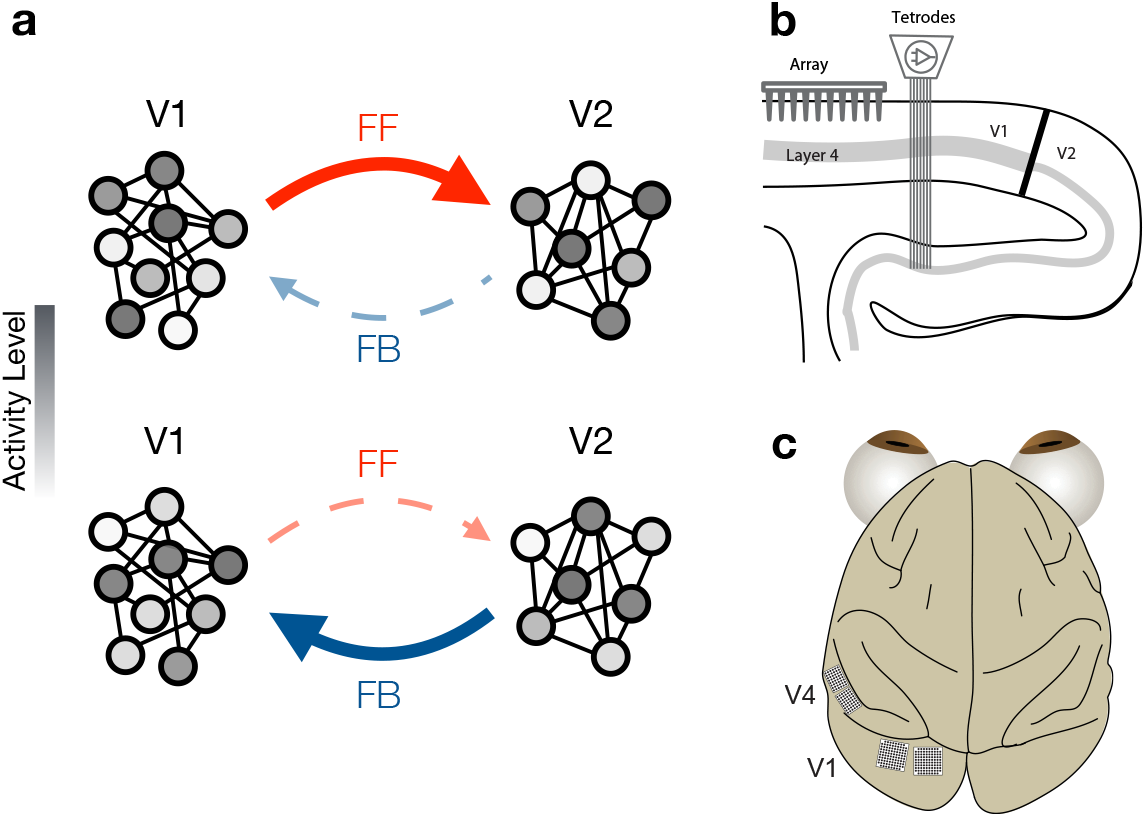
Studying feedforward and feedback interactions using neuronal population activity. **(a)** Each circle represents a neuron in each area, with the shading representing the activity level of the neuron. The population activity patterns involved in feedforward signaling (top) might be distinct from those involved in feedback interactions (bottom). **(b)** Schematic showing a sagittal section of occipital cortex and the recording setup for the V1-V2 recordings. We simultaneously recorded V1 population activity using a 96-channel array and V2 population activity using a set of movable electrodes and tetrodes. **(c)** Schematic showing an overhead view of the recording setup for the V1-V4 awake recordings. We simultaneously recorded V1 and V4 population activity using one 96-channel and one 48-channel array in V1 and a 48-channel array in V4 in the first animal, and two 96-channel arrays in V1 and two 48-channel array in V4 in the second animal.

## Results

We simultaneously recorded from neuronal populations in V1 (88 to 159 neurons; mean: 112.8 ± 12.3 SEM) and V2 (24 to 37 neurons; mean: 29.4 ± 2.4 SEM) in three anesthetized monkeys (Fig. 1b; five recording sessions), as well as in V1 (34 to 128 neurons; mean: 66.6 ± 16.2 SEM) and V4 (12 to 84 neurons; mean: 58.8 ± 12.4 SEM) in two awake fixating monkeys (Fig. 1c; five recording sessions). Animals were shown drifting gratings of different orientations (1280 ms stimulus duration for V1-V2; 200 ms for V1-V4), followed by a blank screen (1500 ms for V1-V2; 150 ms for V1-V4). Recording sites were chosen so that the spatial receptive fields of the V1 and V2/V4 populations overlapped (see ref. 19 and Supplementary Fig. 1).

### Temporal structure of inter-areal interactions

We first characterized the temporal dynamics of the interaction between neuronal population spiking responses in V1 and V2. To do so, we asked: (1) how the interaction evolved during stimulus presentation and the subsequent period of spontaneous activity (which together constitute a trial); and (2) how the interaction depended on the time delay considered between the two areas. Given that these areas are reciprocally connected, with activity flowing in both directions, it is possible that there are periods during which V1 leads V2 activity, and other periods where it lags behind.

To measure interactions between areas, we employed Canonical Correlation Analysis (CCA). Consider representing the activity in two neuronal populations using two activity spaces, one for each area. In each space, each coordinate axis corresponds to the activity of a recorded neuron (Fig. 2a). Within a given time window, the spike counts of the neurons (in the two populations) define a point in each space. For each point in V1 activity space (Fig. 2a, left panel), there is a corresponding, simultaneously recorded point in V2 activity space (Fig. 2a, right panel). CCA seeks dimensions of activity in each area, such that activity along those dimensions is maximally correlated across the two areas (Fig. 2a, bottom panel). For this analysis, we focused on the most correlated dimensions across the two areas (i.e., the first canonical pair; correlations associated with the second canonical pair were on average 60% lower and close to chance level). We used the correlation value for the first canonical pair as a measure of inter-areal interaction strength, which we refer to as *population correlation*.

**Figure 2.**
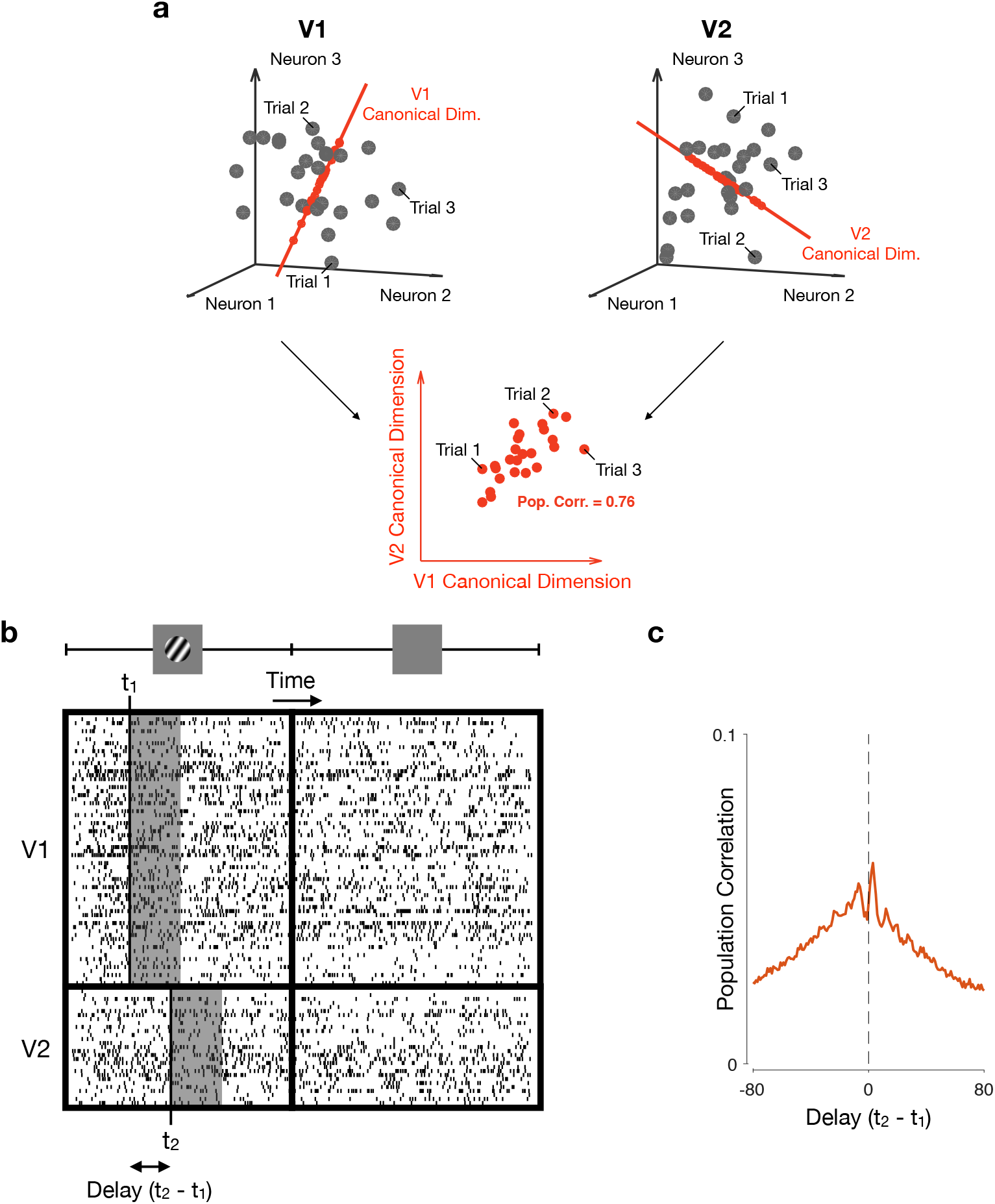
Using Canonical Correlation Analysis (CCA) to capture population interactions. Relating activity across two neuronal populations. Each circle represents the population activity recorded on a given trial. For each activity point observed in the V1 population (left panel; gray dots), there is a corresponding, simultaneously recorded activity point observed in V2 (right panel, gray dots). The red axes represent the first pair of canonical dimensions, identified using CCA. Neuronal activity projected onto the first pair of canonical dimensions (red dots) is highly correlated across the two areas (bottom panel). **(b)** Spike counts across the recorded neurons are taken in specified time windows (gray boxes), which may either be positioned at the same time in both areas (i.e., *t*_H_ = *t*_2_) or with a delay between areas (*t*_1_ 6= *t*_2_). The activity in each gray box is represented by a circle in panel (a). **(c)** The population correlation function corresponds to the correlation between areas returned by CCA (the correlation associated with the first pair of canonical dimensions), as a function of the time delay between areas (*t*_2_ -*t*_1_).

Interactions between areas likely involve time delays due to signal conduction, as well as network processing. This implies that the activity across areas might not be most related for matched (simultaneous) time windows, but for time windows shifted forward or backward in time. Thus, we used CCA to relate activity recorded in V1 with activity in V2 at different time delays (Fig. 2b; Methods) to produce a population correlation function (Fig. 2c). This population correlation function can be computed at different epochs in a trial.

We found that V1-V2 population correlations were lowest just after stimulus onset, increased steadily during stimulus presentation, and were highest for spontaneous activity (Fig. 3a). Focusing on the activity shortly after stimulus onset (“Early Evoked”; 160 ms after stimulus onset), population correlations were larger for positive delays than for negative delays (red trace in Fig. 3b, with peak correlation occuring for a lag of 3 ms), meaning V1 activity was most correlated with V2 activity occurring later in time –consistent with a feedforward interaction. The feedforward interaction became less evident later during the evoked activity period (“Late Evoked”; 1120 ms after stimulus onset; yellow trace in Fig. 3b). After stimulus offset, population correlations were larger for negative delays, so that V2 led V1, suggesting a feedback-dominated interaction (“Spontaneous”, purple trace in Fig. 3b, with a broad peak centered at approximately -15 ms; 2240 ms after stimulus onset). For a more complete characterization, we show in Fig. 3c how population correlations vary as a function of time delay between areas (horizontal axis) and the time relative to stimulus onset (vertical axis; note that the population correlation functions in Fig. 3b represent horizontal slices of this representation).

**Figure 3.**
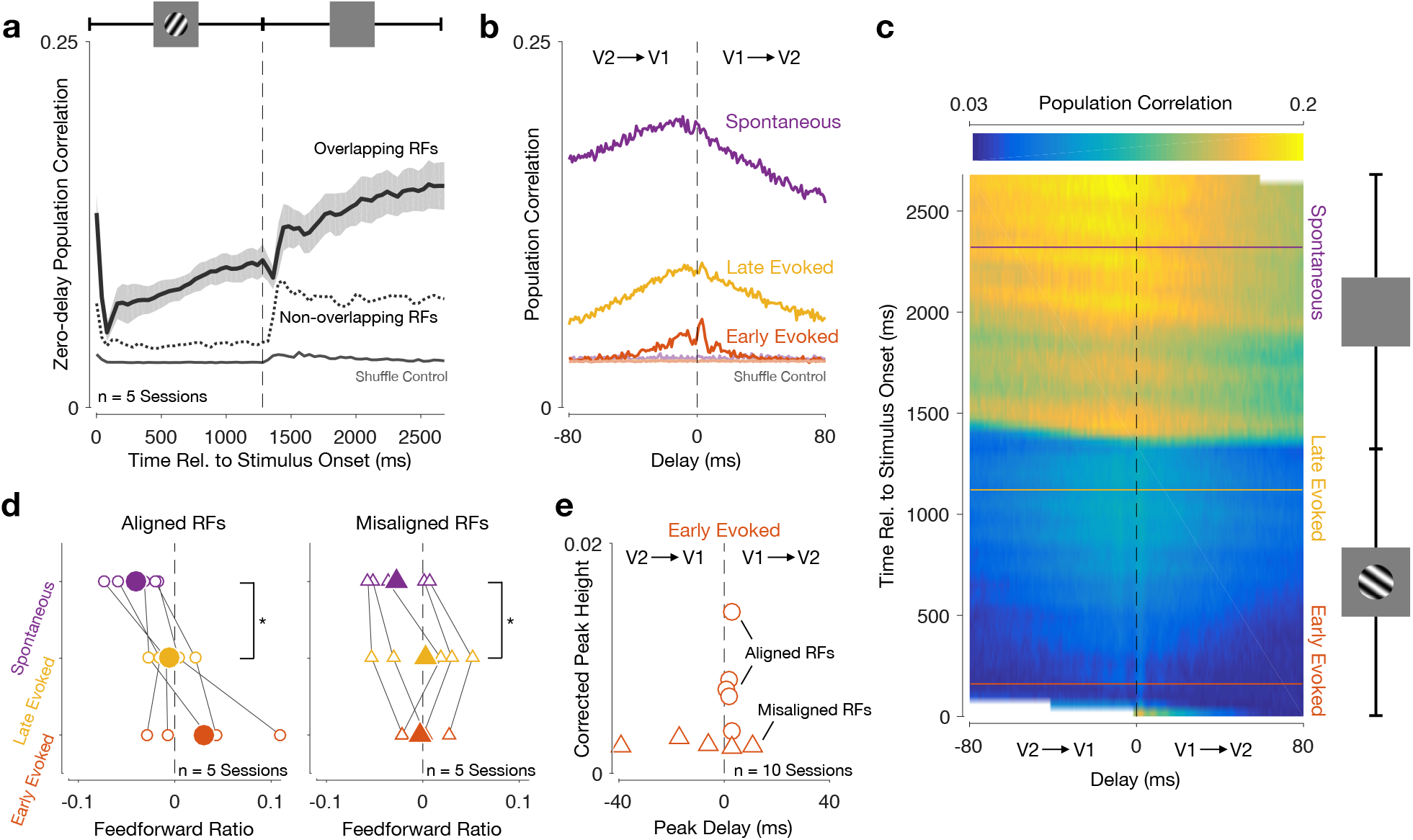
V1-V2 interaction transitions from feedforward-dominated shortly after stimulus onset to feedback-dominated during the spontaneous period. **(a)** Inter-areal zero-delay population correlation increased throughout the trial, and was higher for spontaneous activity than for evoked activity. Zero-delay refers to spike counts taken in the same time window in the two areas (*t*_1_ = *t*_2_ in Fig. 2b). Black line shows the average across all recording sessions for which the V1 and V2 populations have aligned receptive fields. Shading indicates S.E.M. Dotted line shows average across all recording sessions where the the V1 and V2 receptive fields are misaligned. Gray line shows average population correlation after shuffling trial correspondence between the two areas. **(b)** Population correlation functions for an example session (red: early evoked, yellow: late evoked; purple: spontaneous). Faded lines show population correlation functions after shuffling trial correspondence between the two areas (note that there are multiple superimposed lines). **(c)** Population correlations at all times during the trial. The horizontal axis represents the time delay between areas (*t*_1_ -*t*_2_), and the vertical axis represents time relative to stimulus onset (*t*_1_). Horizontal lines (red, yellow, and purple) indicate epochs used in panel (b). Dashed vertical line indicates zero-delay population correlations shown in panel (a). White area denotes times for which population correlations could not be computed: the V2 activity window had reached either the beginning or the end of the trial. Same session as in panel (b). **(d)** Feedforward ratio for different epochs of evoked and spontaneous activity. Left panel shows sessions for which the V1 and V2 populations have aligned receptive fields; right panel shows sessions where the the V1 and V2 receptive fields are misaligned. Solid symbols show the average across all recording sessions, whereas open symbols correspond to each recording session. **(e)** An early feedforward peak is only present in recording sessions where the V1 and V2 populations have aligned receptive fields. Peak height is measured after performing a jitter-correction to isolate fast timescale interactions (see Methods). Circles correspond to recording sessions for which the V1 and V2 populations have aligned receptive fields. Triangles correspond to sessions in which the V1 and V2 receptive fields are misaligned.

To quantify the shift from feedforward-to feedback-dominated interactions, we calculated a feedforward ratio, defined as the difference between the feedforward (positive delay) and feedback (negative delay) sides of the population correlation function, divided by their sum. In every recording session, we found that V1-V2 interactions were more feedback-dominated during the spontaneous period than during the evoked period (Fig. 3d, left; average feedforward ratio, computed in the -80 to 80 ms delay range: -0.005 ± 0.008 SEM for late evoked activity; -0.040 ± 0.011 SEM for spontaneous activity; one-sided paired Wilcoxon signed-rank test, *p* = 0.03 for difference between late evoked and spontaneous activity across all 5 recording sessions; t-test for feedforward ratio, *p* = 0.57 for late evoked activity, *p* = 0.02 for spontaneous activity).

The population correlation functions contain both slow- and fast-timescale features. To isolate the fast-timescale features, particularly evident early in the evoked period (Fig. 3b, red), we computed jitter-corrected population correlation functions for responses measured after stimulus onset^35,36^ and computed their peak location and height (Supplementary Fig. 2). Clear feedforward peaks will have large heights whereas the absence of a peak will result in a small peak height with highly variable peak times (i.e., reflecting “noise” in the correlation function). We found a clear, early feedforward peak in all recording sessions for which the V1 and V2 receptive fields were aligned (Fig. 3e, open circles; average peak height: 0.008 ± 0.002 SEM; average peak delay: 2.2ms ±0.37 SEM).

If the effects shown in Fig. 3 truly reflect feedforward and feedback interactions, they should display appropriate retinotopic specificity. Feedforward connections are more retinotopically precise than feedback connections^37–41^. As a result, feedforward interactions should require retinotopic alignment, whereas feedback interactions might be more tolerant of retinotopic misalignment between the neurons sampled in the two areas. To test this prediction, we performed additional recordings for which the spatial receptive fields of the V1 and V2 populations were misaligned by several degrees (mean center-to-center population spatial receptive field distance was 3.73 deg for misaligned sessions and 0.58 deg for aligned sessions).

Population correlations were lower for these recordings than for those from populations with aligned receptive fields (Fig. 3a, dotted line). The fast time-scale correlation peaks observed shortly after stimulus onset for aligned populations (Fig. 3e, circles) were absent in responses from populations with misaligned receptive fields, evident as small peak heights and inconsistent peak delays (Fig. 3e, triangles; average peak height: 0.0025 ± 0.0001 SEM; one-sided permutation test, *p* = 0.004 for difference between sessions with aligned vs. misaligned receptive fields). Despite the absence of a clear feedforward peak, the V1-V2 interaction for the misaligned populations was still feedback-dominated during spontaneous activity (Fig. 3d, purple; average feedforward ratio: -0.003 ± 0.019 SEM for late evoked activity; -0.027 ± 0.013 SEM for spontaneous activity; one-sided paired Wilcoxon signed-rank test, *p* = 0.03 for difference between late evoked and spontaneous activity across all 5 recording sessions). Thus, the feedforward and feedback interactions identified by CCA have properties consistent with the underlying anatomical specificity.

To test whether the dynamics of V1-V2 interactions might reflect in part changes in the activity within each area, rather than the interaction between areas, we devised two controls. First, we split each V1 and V2 population randomly into two groups, and measured within-area correlations as we had done when analyzing inter-areal interactions. The features described for inter-areal interactions were absent when identical analyses were performed on neurons recorded in the same area (Supplementary Fig. 3). Specifically, within-area interactions showed no evidence of a feedforward peak and were symmetric with respect to the time lag during late evoked and spontaneous activity. Thus, the changes in temporal structure shown in Fig. 3 are specific to inter-areal interactions.

Second, we tested whether the dynamics of inter-areal interactions might be related to differences in neuronal onset latency in the two areas, or to changes in the firing rates over time within each population. To assess this possibility, we performed CCA after shuffling the correspondence of trials in the two areas, while keeping the temporal correspondence within each area intact (see Methods). This shuffling procedure maintained the firing rate time courses and correlation structure within each area, but broke the trial-by-trial correspondence of activity across the two areas. After shuffling, inter-areal correlations no longer increased throughout the trial (Fig. 3a, light trace). Furthermore, there was no evidence of a feedforward interaction early in the trial, nor was there a shift to a feedback-dominated interaction during spontaneous activity (Fig. 3b, light traces). Thus, the dynamics of inter-areal interactions cannot be attributed to different onset latencies or response dynamics in the two areas.

We then asked whether inter-areal interactions showed similar dynamics in responses measured in awake animals as in the responses measured in anesthetized animals considered thus far. We recorded V1 and V4 population activity, in two animals performing a passive fixation task in which drifting gratings were presented (Methods). As with V1-V2 responses, V1-V4 population correlation increased throughout the evoked period (Fig. 4a; compare with Fig. 3a). Just after stimulus onset, V1-V4 interactions were feedforward-dominated (Fig. 4b, red curve; 75 ms after stimulus onset). Notably, the feedforward peak was located at approximately 25 ms delay, longer than the delay of the feedforward peak for the V1-V2 interaction and with a broader profile (compare with Fig. 3b). Over time, the initial feedforward interaction was replaced by a feedback-dominated interaction (Fig. 4b, yellow curve; compare with Fig. 3b; 125 ms after stimulus onset). Figure 4c shows the V1-V4 population correlation functions at all epochs during the trial. The shift from a feedforward-to a feedback-dominated interaction was present for all recording sessions (Fig. 4d; average feedforward ratio, computed in the -50 to 50 ms delay range: 0.088 ± 0.014 SEM for early evoked activity; -0.038 ± 0.008 SEM for late evoked activity; one-sided paired Wilcoxon signed-rank test, *p* = 0.03 for difference between early evoked and late evoked activity across all 5 recording sessions; t-test for feedforward ratio, *p* = 0.020 for early evoked activity, *p* = 0.003 for late evoked activity). Importantly, this temporal structure was absent in interactions between subpopulations within each cortical area (Supplementary Fig. 4), and when we shuffled responses to remove the trial-by-trial correspondence between areas (Fig. 4a,b, faded traces).

**Figure 4.**
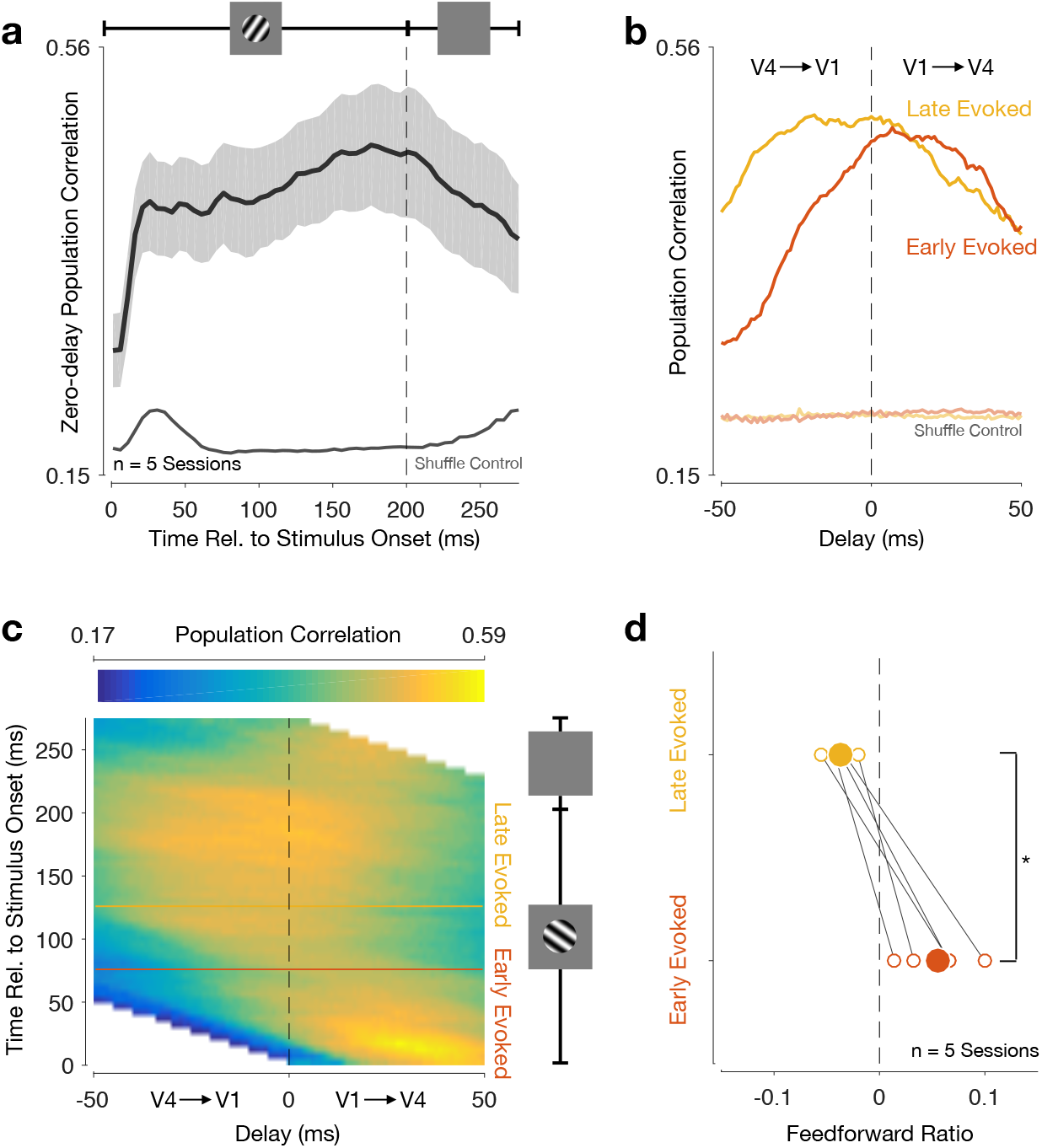
V1-V4 interaction transitions from feedforward-to feedback-dominated during the evoked period. **(a)** Inter-areal zero-delay population correlation increased throughout the evoked period. Black line shows average across all recording sessions. Shading indicates S.E.M. Gray line shows average population correlations after shuffling trial correspondence between the two areas. **(b)** Population correlation functions for an example session, for early (red) and late evoked (yellow) activity. Due to the short duration of the inter-stimulus period, we could not compute a population correlation function for spontaneous activity. Faded lines show population correlation functions after shuffling trial correspondence between the two areas (note that there are multiple superimposed lines). **(c)** Population correlations at all times during the trial. The horizontal axis represents the time delay between areas (*t*_1_ -*t*_2_), and the vertical axis represents time relative to stimulus onset (*t*_1_). Horizontal lines (red and yellow) indicate epochs used in panel (b). Dashed vertical line indicates zero-delay population correlations shown in panel (a). White area denotes times for which population correlations could not be computed: the V4 activity window had reached either the beginning or the end of the trial. Same session as in panel (b). **(d)** Feedforward ratio for early and late evoked activity. Solid circles show the average across all recording sessions, whereas open circles correspond to each recording session.

### Population structure of inter-areal interactions

Past work has suggested that inter-areal interactions are selective, in terms of which population activity patterns are related across areas^29,42^. That is, not all activity fluctuations in one area are reflected in the activity of its downstream targets: some fluctuations remain private to the source area. In our analysis thus far, we have focused solely on the strength and directionality of inter-areal interactions.

Given the observed dynamics of inter-areal interactions, we wondered whether the patterns of activity relayed across areas might be different between feedforward- and feedback-dominated periods. One possibility is that the patterns of activity most related across the two areas are similar during these two periods. Since feedback signaling is hypothesized to alter, or correct, visual representations upstream^43–45^, one might expect that the dimensions most affected by feedback are the same dimensions that are involved in feedforward interactions. This would suggest feedforward and feedback interactions “read from” and “write to” the same population activity patterns, sharing the same communication channel. Alternatively, feedforward and feedback interactions might unfold through separate channels involving distinct population activity patterns, and thus perhaps minimizing how much they directly interact. This would suggest that feedback processing affects dimensions of upstream activity that are not directly involved in relaying visual information downstream.

To distinguish between these possibilities, we divided the trial in epochs and measured how the canonical dimensions identified during one epoch generalized to another. For example, we asked whether the canonical dimensions identified during the feedforward-dominated period (Fig. 5a) captured inter-areal correlations during the feedback-dominated period as well as the canonical dimensions identified during that feedback-dominated period (Fig. 5b). Good generalization would imply that the same patterns of activity were related across areas during periods of feedforward- and feeback-dominated interactions. If, however, the patterns of activity most related across areas differed, the canonical dimensions found during the feedforward-dominated periods would not capture inter-areal correlations during the feedback-dominated periods (Fig. 5c).

**Figure 5.**
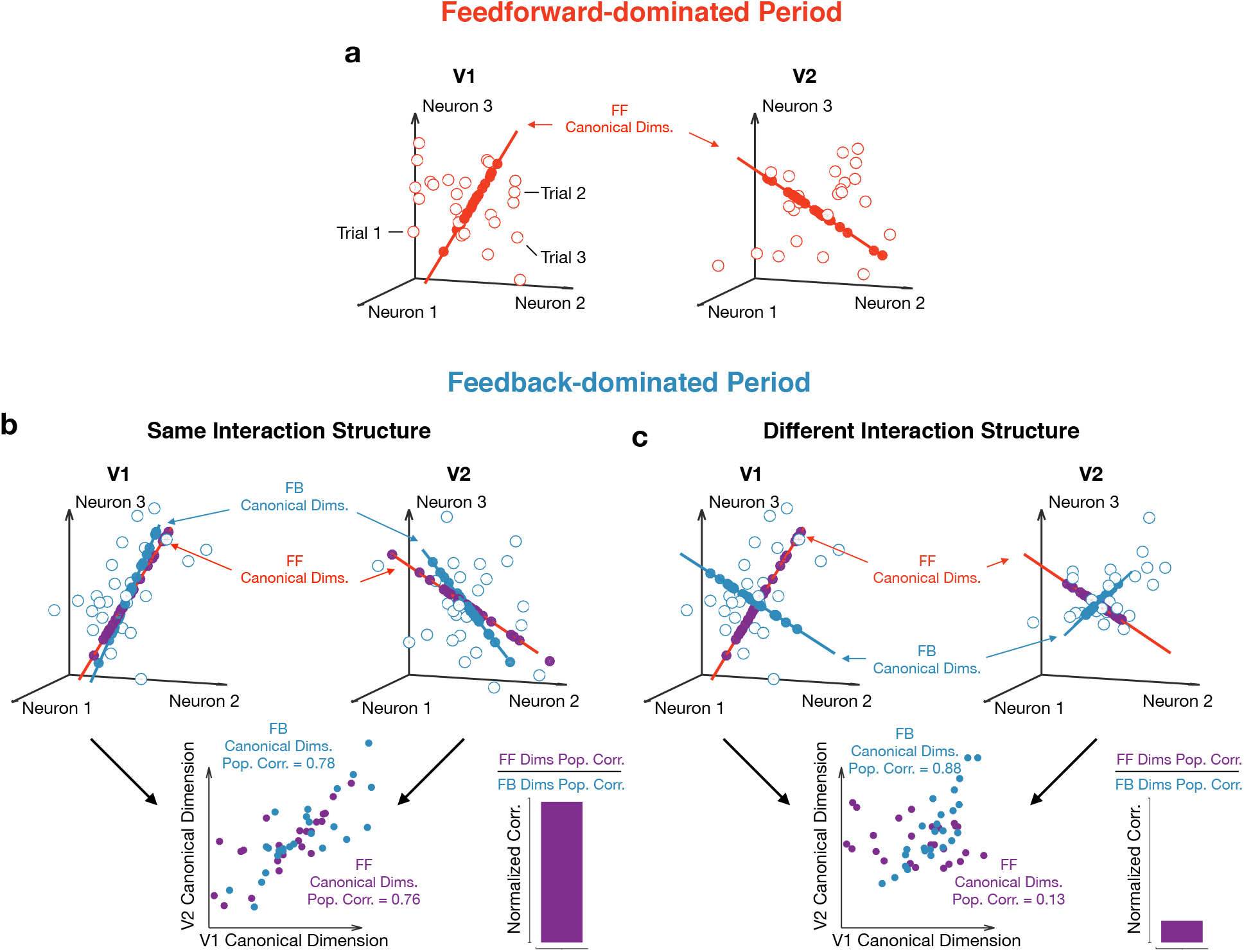
Assessing whether feedforward- and feedback-dominated interactions involve the same population activity patterns. **(a)** Canonical dimensions identified during a feedforward-dominated period in the trial (red dimensions). These are putative “Feedforward” (FF) canonical dimensions. Open red circles denote activity during the feedforward-dominated period. Solid red circles denote the projection onto the FF canonical dimensions. **(b)** We can then ask whether these FF canonical dimensions generalize to a feedback-dominated period. One possibility is that the interaction structure (defined using the canonical dimensions) remains stable across the two periods. In this case, the FF canonical dimensions (red dimensions) capture a similar level of correlation during the feedback-dominated period as the canonical dimensions identified during this period, the putative “Feedback” (FB) canonical dimensions (blue dimensions). As a result, the normalized correlation, the ratio of the population correlation for the FF canonical dimensions to that for the FB canonical dimensions (both computed in a cross validated manner; see Methods), is close to 1. Open blue circles denote activity during the feedback-dominated period. Solid purple circles denote the projection of activity during the feedback-dominated period onto the FF canonical dimensions. Solid blue circles denote the projection onto the FB canonical dimensions. **(c)** Alternatively, the interaction structure might change across the two periods. In this case, the FF dimensions capture only a small fraction of the population correlation during the feedback-dominated period. Same conventions as in panel (b).

We found that dimensions identified early in the evoked activity period, when V1-V2 interactions were feedforward-dominated, did not generalize well to later epochs (Fig. 6a; average normalized correlation, for which a value of 1 indicates perfect generalization: 0.56 ± 0.05 for mid evoked, 0.59 ± 0.04 for late evoked, 0.36 ± 0.04 for late spontaneous; one-sided paired Wilcoxon signed-rank test, *p* = 0.03 for difference between correlation captured using early evoked vs mid evoked, late evoked or late spontaneous dimensions in the corresponding epochs, across all recording sessions). The failure of dimensions identified during the feedforward-dominated period to generalize to the dimensions identified during spontaneous activity suggests that epochs in the feedforward-dominated period involve distinct patterns of population activity compared to epochs in the feedback-dominated period. The generalization was better between epochs later after stimulus onset, when the correlation functions were more symmetric (Fig. 6b; average normalized correlation: 0.64 ± 0.04 for early evoked, 0.94 ± 0.03 for mid evoked, 0.45 ± 0.03 for late spontaneous; one-sided paired Wilcoxon signed-rank test, *p* = 0.03 for difference between correlation captured using early evoked vs mid evoked, late evoked or late spontaneous dimensions in the corresponding epochs, across all recording sessions), indicating that the patterns of activity related between areas are stable for mid and late evoked activity. Dimensions identified during epochs of feedback-dominated interaction, during spontaneous activity, failed to generalize to evoked activity (Fig. 6c; average normalized correlation: 0.47 ± 0.06 for early evoked, 0.50 ± 0.03 for mid evoked, 0.52 ± 0.02 for late evoked; paired one-sided Wilcoxon signed-rank test, *p* = 0.03 for difference between correlation captured using early evoked vs mid evoked, late evoked or late spontaneous dimensions in the corresponding epochs, across all recording sessions). These analyses were carefully designed to focus exclusively on changes in the across-area interaction structure, and to be insensitive to changes in the structure of population activity within each area (see Supplementary Fig. 5, Methods, and Supplementary Information).

**Figure 6.**
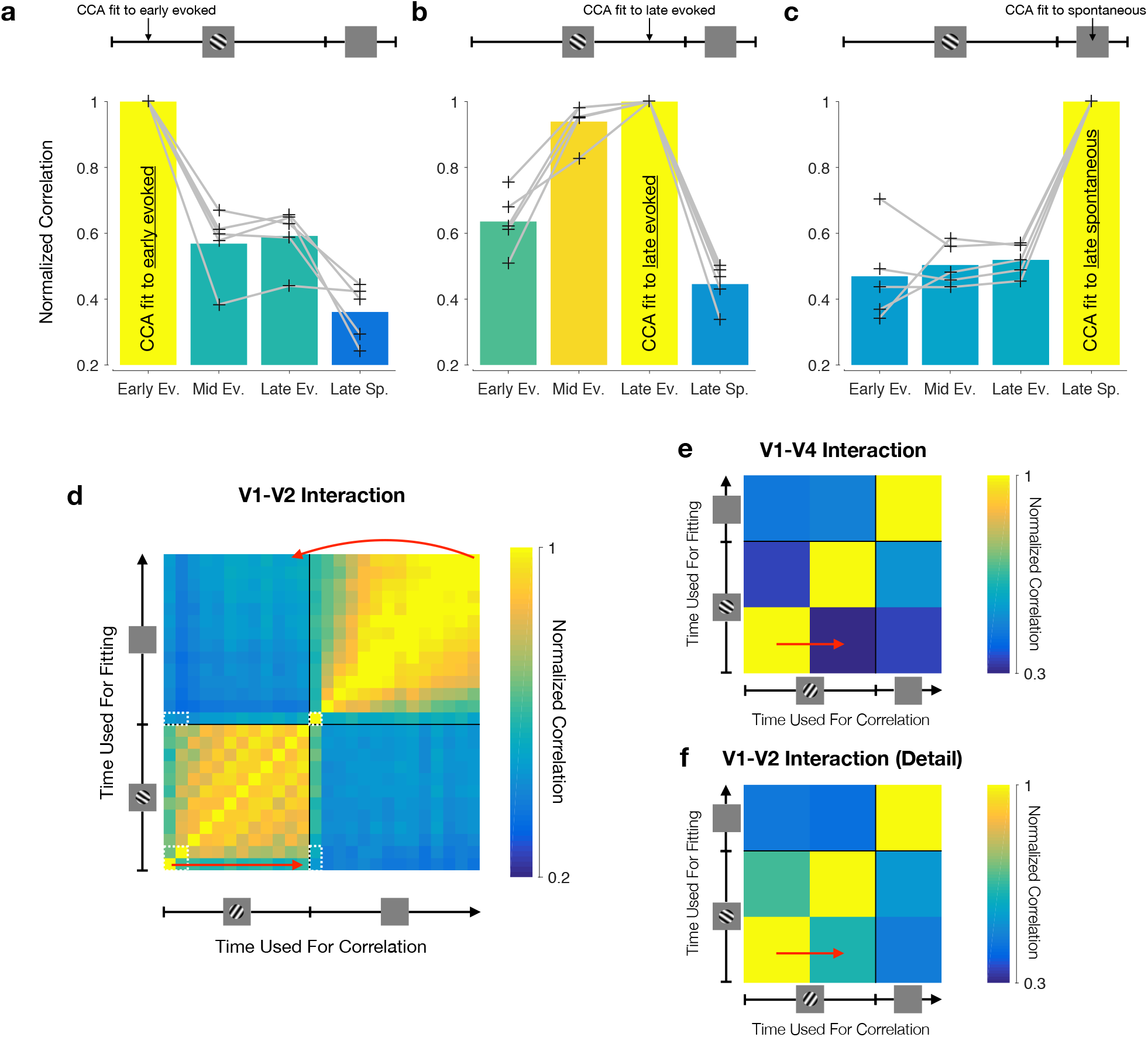
Interaction structure is distinct for the feedforward- and feedback-dominated periods. **(a)** The dimensions found by fitting CCA shortly after stimulus onset (80 ms after stimulus onset) do not generalize well to later epochs in the evoked period, and worse still during the spontaneous period. Grey lines correspond to each of the 5 recording sessions. We report the normalized correlation, defined as the total correlation captured at the test epoch by the dimensions fit to some other epoch over the total correlation captured by the dimensions fit to the test epoch (both computed in a cross-validated manner; see Methods). Dimensions identified late in the evoked period (1180 ms after stimulus onset) do not generalize well to early evoked epochs and to epochs in the spontaneous period, but generalize well to mid-evoked activity. Same conventions as in panel (a). **(c)** Dimensions identified during the spontaneous period do not generalize well to the evoked period. Same conventions as in panel (a). **(d)** Assessing changes in interaction structure across the entire trial. The trial was divided into 100 ms segments, and CCA was applied separately to the activity in each time window. The top two canonical pairs associated with each window were then used to capture inter-areal correlations in the other time windows (see Methods). Each row corresponds to the time during the trial during which the canonical dimensions were identified. Each column corresponds to the time during the trial where the population correlation is assessed. Each entry shows the average across all recording sessions. Straight arrow highlights the comparison of the interaction structure within the evoked period. Curved arrow highlights the comparison of the interactions structure between the spontaneous and the evoked periods. Dashed white boxes indicate epochs reproduced in panel (f). **(e)** Comparing identified dimensions across epochs for the awake V1-V4 recordings. The trial was divided into 100 ms segments, and CCA was applied separately to the activity in each time window. The top canonical pair associated with each window was then used to capture inter-areal correlations in the other time windows (see Methods). Arrow highlights the comparison of the interaction structure within the evoked period. Same conventions as in panel (d). **(f)** Detailed view of the V1-V2 generalization performance for the comparable epochs between the V1-V2 and V1-V4 recordings. Epochs are indicated by the dashed white boxes in panel (d).

To gain a more complete picture, we assessed generalization performance between each possible pairing of epochs for defining canonical dimensions (Fig. 6d, vertical axis), and for testing their relevance (horizontal axis). Each row corresponds to a set of canonical dimensions, identified at a particular epoch, and applied to activity at each of the other epochs. The patterns of generalization performance mirror the changes we observed in the temporal profile of the interaction. As the feedforward interaction weakened after stimulus onset (Fig. 3b, compare red and yellow curves), the patterns of activity most related across the two areas changed as well (Fig. 6d, straight arrow, bottom left). Furthermore, the spontaneous activity period, which was more feedback-dominated than the evoked activity period (Fig. 3d), involved different patterns of activity from those involved in the evoked period (Fig. 6d, curved arrow, top right).

We obtained similar results when analyzing V1-V4 activity. V1-V4 interactions transitioned from a feedforward-to a feedback-dominated interaction (Fig. 4), and the dimensions mediating these interactions changed between these epochs as well (Fig. 6e; the number of epochs is smaller here due to the shorter trial duration; one-sided paired Wilcoxon signed-rank test, *p* = 0.03 for difference between correlation captured using late evoked vs early evoked or late spontaneous dimensions in the corresponding epochs, across all recording sessions). Specifically, the V1-V4 interaction became feedback-dominated at the end of the evoked period (Fig. 4d, yellow circles), and this was accompanied by poor generalization between early and late evoked dimensions (Fig. 6e, straight arrow; average normalized correlation for second epoch of evoked activity for V1-V4: 0.27 ± 0.02). In contrast, the V1-V2 interactions shifted more slowly away from a feedforward-dominated interaction after stimulus onset (Fig. 3d, yellow circles). Consistent with this slower transition, V1-V2 dimensions identified soon after stimulus onset generalized better for nearby epochs of evoked activity, compared to V1-V4 (Fig. 6f, straight arrow; averaged normalized correlation for second epoch of evoked activity for V1-V2: 0.62 ± 0.05).

Taken together, our findings suggest feedforward and feedback inter-areal interactions involve different patterns of population activity. In turn, this implies that the aspects of V1 population activity that are relayed downstream are not necessarily the aspects of activity that are most influenced by feedback. Feedforward and feedback processing might thus occur in separate subspaces of population activity, concurrently and through different “channels”.

## Discussion

We leveraged multi-area recordings to understand the interactions between neuronal population spiking responses in V1 and downstream areas V2 and V4. We found that interactions are feedforward-dominated shortly after stimulus onset, and become feedback-dominated later in the stimulus period and during spontaneous activity. Thus, when a stimulus persists, or when no stimulus is presented, the role of top-down inputs from areas such as V2 and V4 to V1 is more prominent. Furthermore, we found that the population activity patterns most related across areas during feedforward-dominated periods were distinct from those most related during feedback-dominated periods (Fig. 7). This suggests that feedforward and feedback signals involve distinct axes in population activity space, which might allow them to be relayed with minimal direct interference.

**Figure 7.**
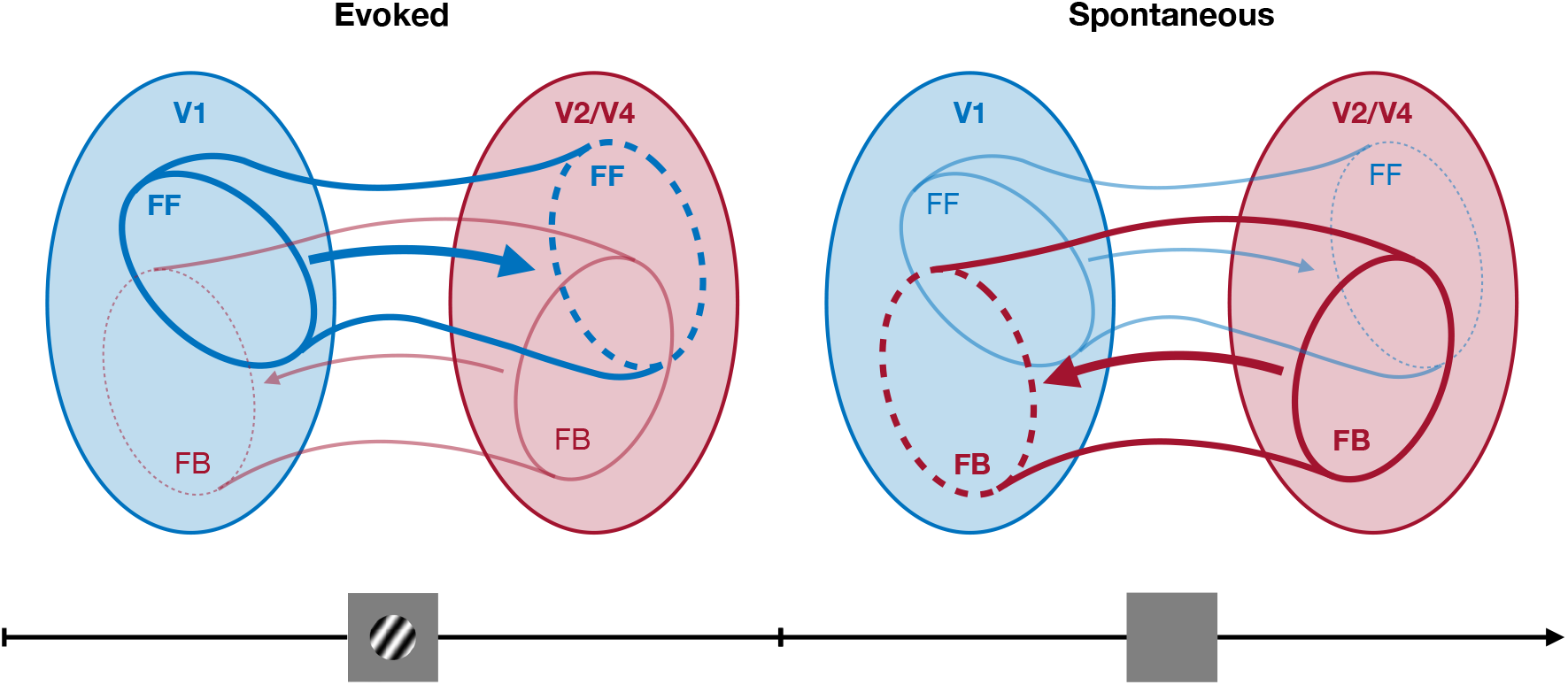
Summary of results. During the early evoked period, interactions between areas tend to be feedforward-dominated. Later during the evoked period and during the spontaneous period, interactions between areas become feedback-dominated. Furthermore, feedforward- and feedback-dominated interactions involve different population activity patterns. Larger ellipses represent the set of all activity patterns one might observe in either the V1 or the V2/V4 populations. The smaller ellipses represent the activity patterns most related across the two areas.

In this study, we measured population correlations in activity across areas at different time lags, and we refer to the identified interactions as feedforward or feedback, based on the lags at which population correlations were maximal. The feedforward interactions that we identified from V1 to V2 are likely to reflect direct (i.e., monosynaptic) input for the following reasons. First, our recordings were performed in the output layers of V1 and input layers of V2^19^. Second, the V1-V2 feedforward peak was sharp, and centered at a delay of 2-3 ms (cf. Fig. 3b,e), consistent with the propagation delay between these areas^46,47^. Third, the feedforward peaks identified for the V1-V2 interactions were absent in recording from neuronal populations with poorly aligned receptive fields (cf. Fig. 3e), consistent with specificity of feedforward connections between these areas^37,38^. In contrast, feedback interactions were less temporally precise than the feedforward interactions, suggestive of a longer signaling loop from V2 back to V1 that may involve polysynaptic paths or shared feedback from more distant areas. These feedback interactions were evident both in recordings from populations with aligned or misaligned receptive fields (cf. Fig. 3d), consistent with the broader visuotopic extent of feedback connections^39–41^. For V1-V4 interactions, both the feedforward and feedback interactions were relatively broad (cf. Fig. 4b), which might be explained by the reduced laminar specificity of our recordings in V4 (chronically implanted arrays, compared to movable tetrodes used in V2), and by a larger number of possible paths by which activity can propagate between these two areas^48,49^.

In both sets of experiments (V1-V2 in anesthetized animals and V1-V4 recordings in awake animals), we observed that interactions were feedforward-dominated shortly after stimulus onset, but this feedforward component subsided, giving way to feedback-dominated interactions. However, there was one notable difference: V1-V2 interactions became feedback-dominated only after the stimulus offset, whereas V1-V4 became feedback-dominated during the late evoked period. This difference could reflect a stronger influence of feedback signaling in the awake state, a difference in the areas involved (V2 vs. V4), or the layers in which the neuronal populations were recorded. That we saw a feedback-dominated interaction at all in the anesthetized recordings might seem surprising, since activity in higher cortical areas, and therefore top-down inputs, might be expected to be diminished by anesthesia. Although it is unclear whether the feedback-dominated interaction we observed is the same as that in an awake animal, we note that V2 is a major source of feedback to V1^48^ and it remains highly responsive under sufentanil anesthesia^12,18,19^.

The transition from feedforward-to feedback-dominated interactions during stimulus drive is broadly consistent with inferences drawn from latency measurements. Because V2 depends on input from V1^50^, one would expect interactions between the areas to be feedforward-dominated immediately after stimulus onset. Spatial contextual effects in V1, which are thought to arise in part from feedback from higher visual areas^4^, are evident 50-100 ms after response onset^11,51,52^, consistent with our observation of a shift away from a feedforward-dominated interaction immediately after response onset to a more balanced (V1-V2) or even feedback-dominated (V1-V4) interaction later in the response. While broadly consistent, our observations significantly extend this prior work. In particular, while measurements of onset may provide information about when feedforward and feedback influences begin, they provide little information about their relative influence once both have been engaged. By using population spiking responses, we are able to see network wide changes in the direction of signaling, as a function of stimulus drive. Our claim that inter-areal interactions switch between being feedforward- or feedback-dominated was not based solely on differences in the time lags at which inter-areal correlations were strongest. It is also supported by our finding that the structure of the population activity that was most correlated between areas was distinct in these different periods. Specifically, we found that the dimensions of population activity that were most related across areas during feedforward signaling periods were distinct from those that were most related during feedback periods. The relevant activity patterns were highly reliable: during spontaneous activity or the sustained epochs of evoked activity, the dimensions of activity that were most correlated across areas were consistent in time. Yet, when networks switched from feedforward to feedback signaling (or vice-versa), the relevant activity patterns changed abruptly.

Determining the population structure of inter-areal interactions requires great care. In particular, it is important to ensure that apparent changes in inter-areal interactions do not arise solely from changes in the structure of activity within each area (see ref. 53, in press). For instance, a change in activity structure within one area might cause the canonical dimensions identified to change, even if the manner in which activity in the two areas is related is unchanged (see Supplementary Information for an extended discussion). To avoid such confounds, we defined interaction structure using across-area covariance, and measured changes in this structure so as to only reflect changes in the activity subspaces in each area spanned by the across-area covariance. In addition, we confirmed that our approach did not detect interaction changes when the across-area covariance was held fixed (Supplementary Fig. 5).

In previous work, we reported that V1 interactions with V2 occur through a communication subspace, which defines which population activity patterns are related across areas^29^. The communication subspace was identified using reduced rank regression (RRR), a dimensionality reduction technique related to CCA but different in its technical details (for a review, see ref. 53, in press). Here we chose to use CCA because it treats the population activity of each area symmetrically. This allows us to study feedforward and feedback influences using the same analysis. In contrast, RRR treats each population differently – one area is labeled the “source” (the independent variable in linear regression) and the other area is labeled the “target” (the dependent variable). Although RRR and CCA need not identify the same dimensions, we found that a communication subspace was also evident when employing CCA. Namely, a smaller number of canonical dimensions was required to capture across area correlations compared to within-area correlations (Supplementary Fig. 6).

How do our observations of feedforward and feedback interactions inform our understanding of how these forms of signaling contribute to cortical function? While the computational role of feedforward signaling has been extensively investigated, the role of feedback is more enigmatic. Feedback signals have been proposed to improve or correct feedforward signals, e.g., by providing prior information about the sensory input^43–45^, by providing a prediction of that input (in predictive coding)^54–56^, or by signaling deviations from some higher-order “teaching” signal (in biologically plausible backpropagation)^57–59^. We find that inter-areal interactions just after stimulus onset are feedforward. This might be explained by the abrupt transition from one visual environment to another when a stimulus suddenly appears. Assuming the trial structure is not learned by the visual cortex, stimulus onset is unpredicted or unexpected; according to predictive coding principles, such input should give rise to potent feedforward signaling. As the stimulus persists, inter-areal interactions become feedback-dominated. This transition might indicate that higher cortex is providing signals that attempt to ‘explain away’ the constant, persistent visual input, and thereby reduce responsivity in lower cortex. Interactions are also feedback-dominated during spontaneous activity. This finding is consistent with proposals that sensory representations combine prior information from higher cortex with sensory drive from the periphery. In the absence of overt visual input (i.e., during spontaneous activity), one would expect responses to reflect more strongly the prior, which would be evident as a top-down dominant interaction.

Our finding that feedforward and feedback interactions involve different patterns of population activity may offer a solution to a central enigma in proposals of how feedback contributes to sensory processing: feedback that is too weak may fail to properly modify representations of the sensory stimulus, but feedback that is too strong may contaminate the representation and lead to hallucinations. One solution for providing robust feedback but allowing some flexibility in how it interacts with the bottom-up sensory representation could be to have these occupy different dimensions of V1 population activity, as we find. The presence of the feedback signal in a target area can then be decoupled from the strength of its influence. This would suggest that the balance between feedforward and feedback signaling in sensory cortex might be achieved using the same principles used by motor cortex to generate preparatory signals without causing muscle contractions^42^, by prefrontal networks that host competing sensory inputs but can flexibly switch which one drives the local activity^60^, or by visual cortical areas to selectively communicate^29^.

## Acknowledgements

We thank E. Gokcen and A. Motiwala for invaluable discussions and for providing feedback on the manuscript. This work was supported by the Fundação para a Ciência e a Tecnologia graduate scholarship SFRH/BD/52069/2012 (J.D.S.), John and Claire Bertucci Graduate Fellowship (J.D.S.), NIH U01 NS094288 (C.K.M.), Simons Collaboration on the Global Brain 364994 (B.M.Y., A.K.), 543009 (C.K.M.), 543065 (B.M.Y.), 542999 (A.K.), NIH R01 HD071686 (B.M.Y.), NIH CRCNS R01 NS105318 (B.M.Y.), NIH CRCNS R01 MH118929 (B.M.Y.), NIH R01 EB026953 (B.M.Y.), NSF NCS BCS 1533672 and 1734916 (B.M.Y.), NIH EY016774 (A.K.).

## Author Contributions

J.D.S., C.K.M., A.K. and B.M.Y. designed the analyses. J.D.S. performed all the analyses. A.I.J., A.Z., A.A. and A.K. designed and performed the experiments. J.D.S., C.K.M., A.K. and B.M.Y. wrote the manuscript. C.K.M., A.K. and B.M.Y. contributed equally to this work.

## Methods

### Recordings and visual stimulation

#### Anesthetized V1-V2

Animal procedures and recording details have been described in previous work^19,36^. Briefly, animals (macaca fascicularis, male, 2-3 years old) were anesthetized with ketamine (10 mg/kg) and maintained on isoflurane (1%-2%) during surgery. Recordings were performed under sufentanil (typically 6-18 mg/kg/hr) anesthesia. Vecuronium bromide (150 mg/kg/hr) was used to prevent eye movements. The duration of each experiment (which comprised multiple recording sessions) varied from 5 to 7 days. All procedures were approved by the IACUC of the Albert Einstein College of Medicine.

The data analyzed here are those reported in ref. 29, and a subset of recording sessions reported in ref. 19. Activity in V1 was recorded using a 96 channel Utah array (400 micron inter-electrode spacing, 1 mm length, inserted to a nominal depth of 600 microns; Blackrock, UT). We recorded V2 activity using a set of electrodes/tetrodes (interelectrode spacing 300 microns) whose depth could be controlled independently (Thomas Recording, Germany). These electrodes were lowered through V1, the underlying white matter, and then into V2. Within V2, we targeted neurons in the input layers. We verified the recordings were performed in the input layers using measurements of the depth in V2 cortex, histological confirmation (in a subset of recordings), and correlation measurements. For complete details see ref. 19. Voltage snippets that exceeded a user-defined threshold were digitized and sorted offline. The sampled neurons had spatial receptive fields within 2-4 deg of the fovea, in the lower visual field.

We measured responses evoked by drifting sinusoidal gratings (1 cyc/deg; drift rate of 3-6.25 Hz; 2.6-4.9 deg in diameter; full contrast, defined as Michelson contrast, (*L*_*max*_ - *L*_*min*_)*/*(*L*_*max*_ + *L*_*min*_), where *L*_*min*_ is 0 cd/m^2^ and *L*_*max*_ is 80 cd/m^2^) at 8 different orientations (22.5 deg steps), on a calibrated CRT monitor placed 110 cm from the animal (1024 x 768 pixel resolution at a 100 Hz refresh rate; EXPO). Each stimulus was presented 400 times for 1.28 s. Each presentation was followed by an interval of 1.5 s during which a gray screen was presented.

We recorded neuronal activity in three animals. In two of the animals, we recorded in two different but nearby locations in V2, providing distinct middle-layer populations, yielding a total of five recording sessions.

#### Awake V1-V4

Animal procedures and methods have been reported previously in previous work^61^. In brief, animals (two male, adult cynomolgus macaques) were trained to maintain fixation on a small spot (0.2 × 0.2 deg, 80 cd/m2) on a gray background (40 cd/m2) within a 1.08-1.4 degree diameter fixation window. Eye-position was monitored using a video tracking system (Eyelink II, SR research, ON, Canada) with a sampling rate of 500 Hz. Stimuli were presented on a calibrated monitor 64 cm away from the animal (1024 x 768 resolution for monkey 1, 1400×1050 for monkey 2; 100 Hz refresh rate). After training, Utah arrays (0.4 mm spacing; 1 mm electrode length, Blackrock, UT) were implanted in V1 and V4. For monkey 1 we implanted one 96 channel and one 48 channel array in V1 and one 48 channel array in V4. Monkey 2 had two 96 channel arrays in V1 and two 48 channel arrays in V4 (see Fig. 1c). We targeted the arrays to have matching retinotopic locations in V1 and V4 by relying on anatomical markers and previous mapping studies. Receptive fields were in the lower right visual hemifield and largely overlapping for V1 and V4 populations in both monkeys (Supplementary Fig. 1). All procedures were approved by the IACUC of the Albert Einstein College of Medicine.

Extracellular voltage signals were amplified and band-pass filtered between 250 and 7.5 kHz using commercial acquisition software (Blackrock Microsystems, UT and Grapevine, Ripple, UT). Voltage snippets that exceeded a user-defined threshold were digitized and sorted offline.

Visual stimuli and task contingencies were presented using custom openGL software (EXPO). We used full-contrast sinusoidal drifting gratings (spatial frequency 2 cyc/deg; drift rate: 5 Hz). Stimulus position and diameter were chosen to maximize visual responses. Stimulus diameter was set to 2.5 deg for monkey 1 and 7 deg for monkey 2. Each recording session involved four grating orientations, chosen such that there were two pairs of orientations 5 deg apart, and 90 deg between the two pairs (e.g., 0, 5, 90, 95 deg).

Trials began with the animal fixating on a small spot in the center of the screen. After a delay of 100 ms we presented a random series of gratings (three for monkey 1, four for monkey 2). Each stimulus presentation lasted for 200 ms and was followed by an inter-stimulus interval of 150 ms (grey screen). Animals were positively reinforced with a liquid reward if fixation was maintained throughout the trial. Animals performed on average 1080 ± 255 trials, resulting in 3721 ± 1081 stimulus presentations per session. We recorded neural activity for three sessions in monkey 1 and two sessions in monkey 2.

### Data preprocessing

#### Anesthetized V1-V2

In order to capture how moment-to-moment fluctuations in spiking activity were related across the two areas, we subtracted the corresponding peri-stimulus time histogram (PSTH) from each spike train, which was computed separately for each neuron and grating orientation (after z-scoring the activity of each neuron separately for each of the 8 grating orientations). The PSTH was computed across the entire trial period, including the stimulus presentation period and the subsequent inter-trial period. The resulting residual activity was then pooled across all 8 grating orientations for each recording session. These residual fluctuations can be interpreted as perturbations of the “signal”, or mean activity across trials. By focusing on perturbations of the signal, we can then use linear methods such as CCA (see below) as a local linear approximation to what is likely a globally non-linear relationship of activity across areas ^29,62^. For all analyses, we excluded neurons that fired less than 0.5 spikes/s on average across all trials.

#### Awake V1-V4

To minimize the influence of adaptation effects, we analyzed activity across only the second and third grating presentations, for which V1-V4 responses were qualitatively similar (and smaller than the response to the first stimulus presentation). Activity for each neuron was z-scored separately for the second and third grating presentations, and for each of the 4 grating orientations. As with the V1-V2 recordings, we subtracted the corresponding PSTH from each trial, which was computed separately for each neuron and stimulus condition (i.e., combination of grating orientations). The PSTH was computed across the entire trial period, including the stimulus presentation period and the subsequent inter-trial period. The resulting residual activity was then pooled across all stimulus conditions for each recording session. We observed cross-talk between a small proportion of electrode pairs (average across recording sessions: 1.3% ± 0.8% SEM), evident as a surfeit (*>* 0.025 coincidences/spike) of precise (0.1 ms) synchronous events. We addressed this by removing one of the electrodes in each affected pair. For all analyses, we excluded neurons that fired less than 0.5 spikes/s on average, across all trials.

#### Population correlation functions

When computing the population correlation functions for the anesthetized V1-V2 recordings (Fig. 3), we sought to focus on fast time-scale interaction effects. For this reason, we counted spikes in 1 ms non-overlapping bins. For the awake V1-V4 recordings (Fig. 4), due to the smaller number of trials per recording session and the longer conduction delay between V1 and V4^7^ we counted spikes in non-overlapping 25 ms bins.

#### Interaction structure analysis

For the interaction structure analysis (Fig. 6), for which we were interested in estimating the activity patterns most correlated across areas, we counted spikes in 100 ms non-overlapping bins. Activity was binned starting 50 ms after stimulus onset and extending until the end of the stimulus presentation period (1.2 s of evoked activity) and then starting 50 ms after stimulus offset and extending until the end of the inter-trial period (1.45s of spontaneous activity). We used larger time bins than for computing population correlation functions to increase the reliability of the estimated population activity patterns, in exchange for less temporal resolution. Likewise, for the awake V1-V4 recordings we counted spikes in 100 ms non-overlapping bins, starting 50 ms after stimulus onset and extending until the end of the trial (150 ms of evoked activity and 150 ms of spontaneous activity, for a total of 300 ms).

### Population correlation analysis

In order to capture population correlations between cortical areas, we used Canonical Correlation Analysis (CCA)^63^. CCA finds pairs of dimensions, one in each area, such that the correlation between the projected activity onto these dimensions is maximally correlated:

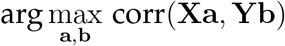

where X is a *n* × *p*_*x*_ matrix containing the residual activity in the V1 population, Y is a *n* × *p*_*y*_ matrix containing the residual activity in the V2 (or V4) population, *n* represents the number of data points, and *p*_*x*_ and *p*_*y*_ are the number of recorded neurons in each of the two areas, respectively. The vectors a and b have dimensions *p*_*x*_ × 1 and *p*_*y*_ × 1, respectively defining dimensions in the population activity space of each area. CCA can find additional pairs of dimensions, by requiring that subsequent pairs are uncorrelated with those previously identified.

In order to measure population correlations at different epochs in the trial, and at different time delays between the areas, we defined two windows of activity, one in each area. Window length was 80 ms for the V1-V2 recordings, and 75 ms for the V1-V4 recordings. Activity was then binned inside each window using 1 ms bins for the V1-V2 recordings (80 data points per window), and 25 ms for the V1-V4 recordings (3 data points per window). The reported results were robust to the specific binning and window length chosen, over a reasonable range.

CCA was then applied to the residual activity taken from all trials within these windows. Given two windows of activity starting at times *t*_1_ and *t*_2_ (relative to the start of the trial), 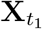 and 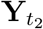, the population correlation between the two areas is given by:

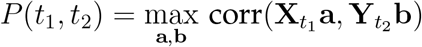

Defining the time within the trial as *t* = *t*_1_ and the delay between the activity in the two areas as *d* = *t*_2_ - *t*_1_, each entry in the population correlation function is given by:

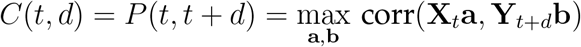

CCA tended to identify only one pair of dimensions with highly significant population correlations: correlations associated with the second canonical pair were on average 60% lower than for the first pair and close to chance level. As such, we constructed the population correlation functions using the first pair of canonical dimensions.

To isolate fast-timescale features in the early evoked activity (Fig. 3e), we computed jitter-corrected population correlation functions. To do so, we jittered the spike times (25 ms jitter window) following the procedure in ref. 36. We then computed population correlation functions using 1 ms binning and a window length of 480 ms, starting 80 ms after stimulus onset, for both the residual activity and the jittered activity. Finally, we subtracted the jittered population correlation function from the population correlation function based on the residual activity, obtaining the jitter-corrected population correlation function. Corrected peak height and delay was computed by finding the maximum of the jitter-corrected population correlation function, as well as the corresponding delay. Supplementary Fig. 2 illustrates this process.

### Comparing interaction structure across time

To determine whether the population activity patterns involved in inter-areal interactions changed during the trial, we leveraged the probabilistic extension of CCA (pCCA)^64^. pCCA is closely related to CCA in that both methods identify the same canonical dimensions. The advantage of pCCA is that it defines an explicit generative model, which we can leverage for model comparison and selection (see Supplementary Information).

Note that the population correlation functions described above could have been computed using pCCA instead of CCA, which would have yielded the same results. We focused there on the first canonical dimension, and did not need the model comparison and selection procedures described below. Thus, solely for clarity of presentation, we opted to introduce the population correlation functions using CCA.

pCCA is defined by the following generative model:

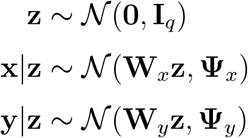

where z is a *q* × 1 latent variable, x and y correspond to the neuronal activity recorded in each of two cortical areas, with dimensionalities *p*_*x*_ × 1 and *p*_*y*_ × 1, respectively, *p*_*x*_ and *p*_*y*_ are the number of neurons recorded in each area, and *q* ≤min(*p*_*x*_, *p*_*y*_). The identity matrix **I**_*q*_ has dimensions *q* × *q*. The mapping matrices **W**_*x*_ and **W**_*y*_ have dimensions *p*_*x*_ × *q* and *p*_*y*_ × *q*, respectively. The covariance matrices **Ψ** _*x*_ and **Ψ** _*y*_ have dimensions *p*_*x*_ × *p*_*x*_ and *p*_*y*_ × *p*_*y*_, respectively. We assume, without loss of generality, that x and y are mean-centered. To fit pCCA, we first applied CCA and used the canonical dimensions and associated canonical correlations to compute the parameters of the pCCA model (see Supplementary Information).

Under the pCCA model, the inter-areal covariance is fully determined by the matrices **W**_*x*_ and **W**_*y*_ (see Supplementary Information for an extended discussion of pCCA and its relation to classical CCA). In particular, the column spaces of these matrices define the activity patterns, in each area, along which activity covaries across the two populations.

We used pCCA to compute the **W**_*x*_ and **W**_*y*_ matrices at different epochs and compared these matrices, across epochs, to assess whether similar population activity patterns were involved in the inter-areal interaction (Fig. 6). We computed the population activity patterns related across areas (i.e., **W**_*x*_ and **W**_*y*_) at one epoch in the trial, and asked how much inter-areal correlation these population activity patterns explained at a different epoch.

Specifically, we first fit a pCCA model with dimensionality *q* (see procedure below for selecting *q*) separately for each epoch *t*, yielding parameters 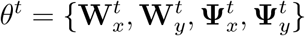. We then asked: given the observed (sample) within-area covariance matrices at time *t*, 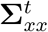 and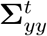, how correlated would the activity across the two areas be if instead of the estimated matrices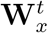 and 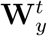, the interaction was instead described by the matrices 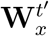 and 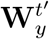, obtained from a different epoch *t*′? In other words, how much does the across area correlation change if we compute across-area correlations using population activity patterns defined by 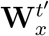 and 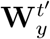, instead of 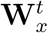 and 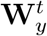? To quantify the change in correlation, we computed normalized correlations, defined as the total correlation captured at epoch *t* by the dimensions fit to epoch *t* ′ over the total correlation captured by the dimensions fit to epoch *t* (both computed in a cross-validated manner; see Methods). Misalignment between the column spaces will lead to decreased correlations, and low normalized correlation. On the other hand, if the mapping matrices 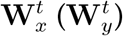 and 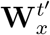 (resp. 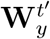)share the same column space (i.e., if the across-area correlations at epochs *t* and *t* ′ involve the same population activity patterns), the resulting correlations should remain the same, and normalized correlation will close to 1. Algorithm (1) describes this procedure in detail (Supplementary Information).

In order to combine results across recording sessions, in Fig. 6 we used a single value for the latent dimensionality *q* for all sessions. To select the value of the latent dimensionality *q*, we first determined the value *q*^*t*^ that maximized the cross-validated data likelihood for each epoch *t*, in each recording session. For the anesthetized V1-V2 recordings, the average dimensionality across all recording sessions was 3.30 ± 0.09 SEM across epochs in the evoked period and 2.09 ± 0.14 SEM across epochs in the spontaneous period (averages taken across epochs and recording sessions). To avoid comparing spurious canonical dimensions, we choose *q* to be no greater than both these estimated dimensionalities. Thus, we choose *q* =2 for these recordings. For the awake V1-V4 recordings the average dimensionality across all recording sessions was 1.8 ± 0.29 SEM across epochs in the evoked period and 2.40 ± 0.24 SEM across epochs in the spontaneous period (averages taken across epochs and recording sessions). Thus, we choose *q* =1 for these recordings. For both sets of recordings, results were robust to different choices of *q*, over a reasonable range.

## Data availability

V1-V2 data are available at the CRCNS data sharing web site, at https://doi.org/10.6080/K0B27SHN. V1-V4 data will be made available upon reasonable request.

## Code availability

MATLAB code that supports the data analyses will be made publicly available upon publication.

---

**Supplementary Figure 1.**
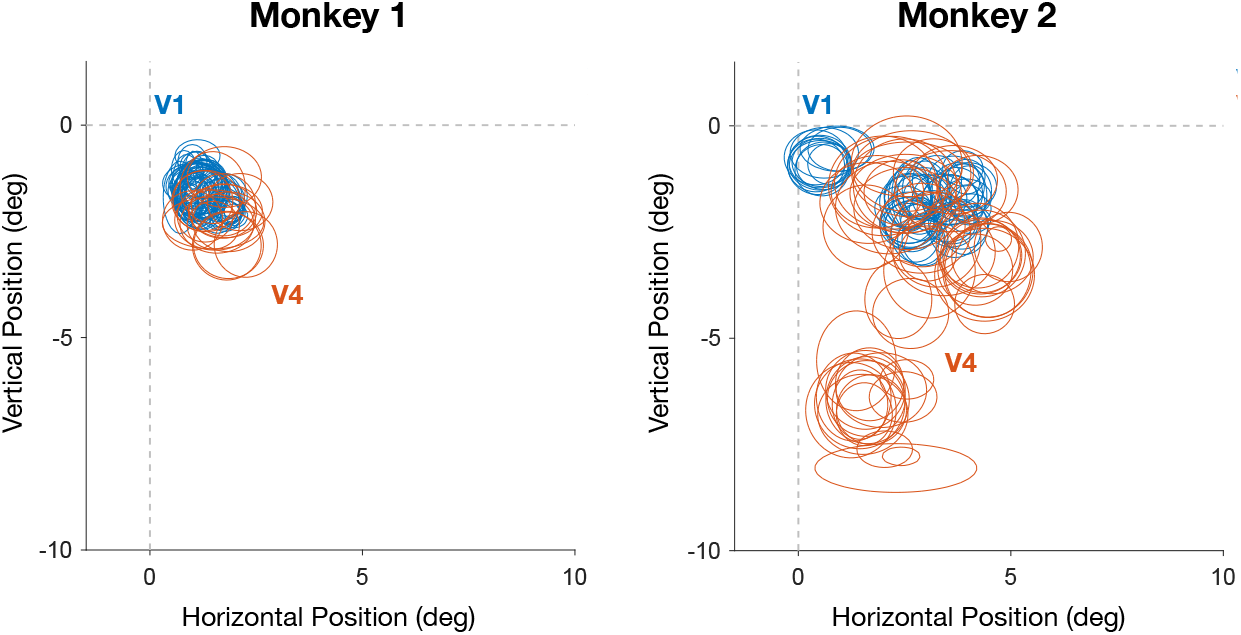
Spatial receptive fields for the V1-V4-awake recordings. Lines indicate 60% contour lines of a 2-dimensional Gaussian fit to the receptive fields. Receptive fields were fitted to unsorted multi-unit activity recorded on each channel.

**Supplementary Figure 2.**
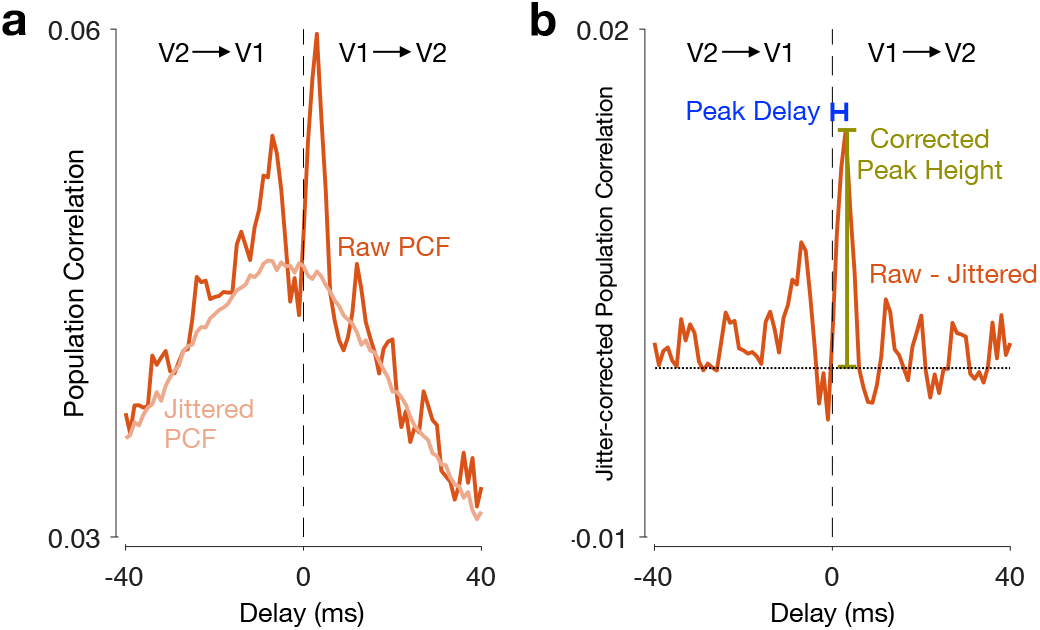
Isolanting feedforward peaks using a jitter-corrected population correlation function. **(a)** If a feedforward peak is caused by precise spiking coordination across the two areas, it should still be present after the slow-timescale component of the population correlation function is removed. To remove the slow-timescale component, thereby isolating fast-timescale features in the early evoked activity, we computed a jitter-corrected population function^36^. We first computed a jittered population correlation function (Jittered PCF; 25 ms jitter window), as described in ref. 36. We then obtained a jitter-corrected PCF by subtracting the jittered PCF from the PCF based on residual activity (Raw PCF). **(b)** We computed the peak height by finding the maximum value of the jitter-corrected PCF, as well as the corresponding time delay. In this session, a clear peak can be observed at 3 ms. Results across all recording sessions are shown in Fig. 3e.

**Supplementary Figure 3.**
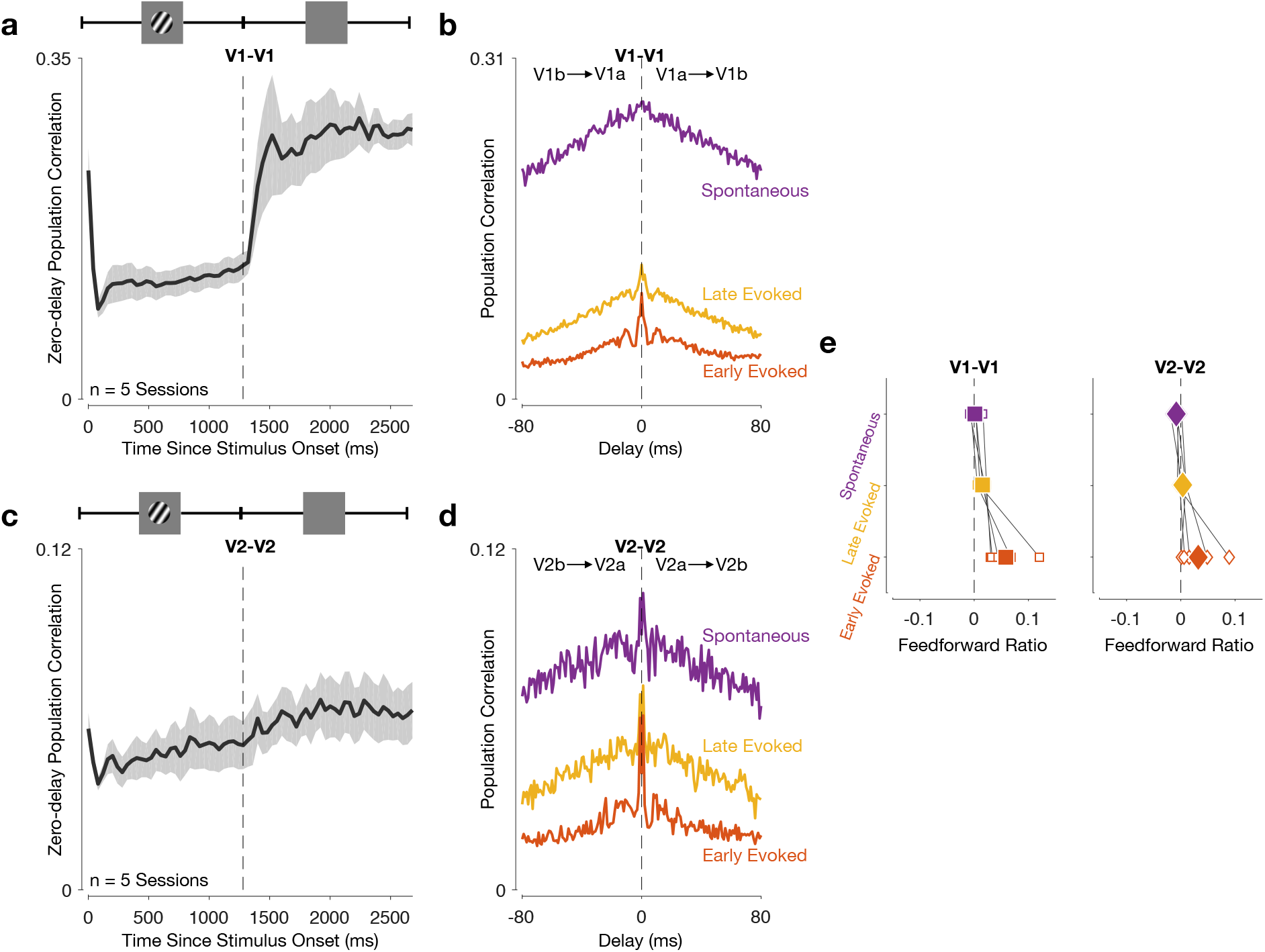
Feedforward peak is absent, and feedback-dominated interactions are not evident between subpopulations within V1 or V2. To test whether the effects in Fig. 3 were specific to inter-areal interactions, we randomly divided the neurons in each area into two subpopulations and computed population correlation functions between the subpopulations for each area. **(a)** V1-V1 zero-delay population correlation increases throughout the trial, and is higher for spontaneous activity than for evoked activity. Solid line shows average across all recording sessions. Shading indicates S.E.M. V1-V1 population correlation functions for an example session (taken as the average across 10 random divisions into two V1 subpopulations). Same conventions as in Fig. 3b. V2-V2 zero-delay population correlation increases throughout the trial, and is higher for spontaneous activity than for evoked activity. Solid line shows average across all recording sessions. Shading indicates S.E.M. **(d)** V2-V2 population correlation function for an example session (taken as the average across 10 random divisions into two V2 subpopulations). Same conventions as in Fig. 3b. **(e)** There are two key features of inter-areal interactions revealed in Fig. 3 which are not present for within-area interactions. First, the feedforward peaks of the population correlation functions for within-area interactions are centered at 0 ms delay (see panels b and d). This is in contrast to the across-area (V1-V2) case, where there is a feedforward peak shortly after stimulus onset (Fig. 3b,e). Second, within-area interactions (V1-V1, left panel; V2-V2, right panel) were neither feedforward-nor feedback-dominated during spontaneous activity (average spontaneous activity feedforward ratio, computed in the -80 to 80 ms delay range: 0.002 ± 0.004 SEM for V1-V1; -0.007 ± 0.003 SEM for V2-V2; t-test for spontaneous activity feedforward ratio, *p* = 0.71 for V1-V1; *p* = 0.09 for V2-V2. This is in contrast to the across-area (V1-V2) case, where interactions were feedback-dominated during spontaneous activity (Fig. 3d). Note that the feedforward ratio is slightly positive for the early evoked period, although the population correlation functions peak at 0 ms time delay throughout the whole trial. This reflects the slightly greater area under the right half compared to the left half of the population correlation function (panels b and d), likely due to the strong change in correlations at stimulus onset (panels a and c). This effect occurs on a slow timescale and motivates our use of jitter-corrected responses reported in the main text (Fig. 3e). Solid symbols show average across all recording sessions, empty symbols correspond to each recording session. Same conventions as in Fig. 3d.

**Supplementary Figure 4.**
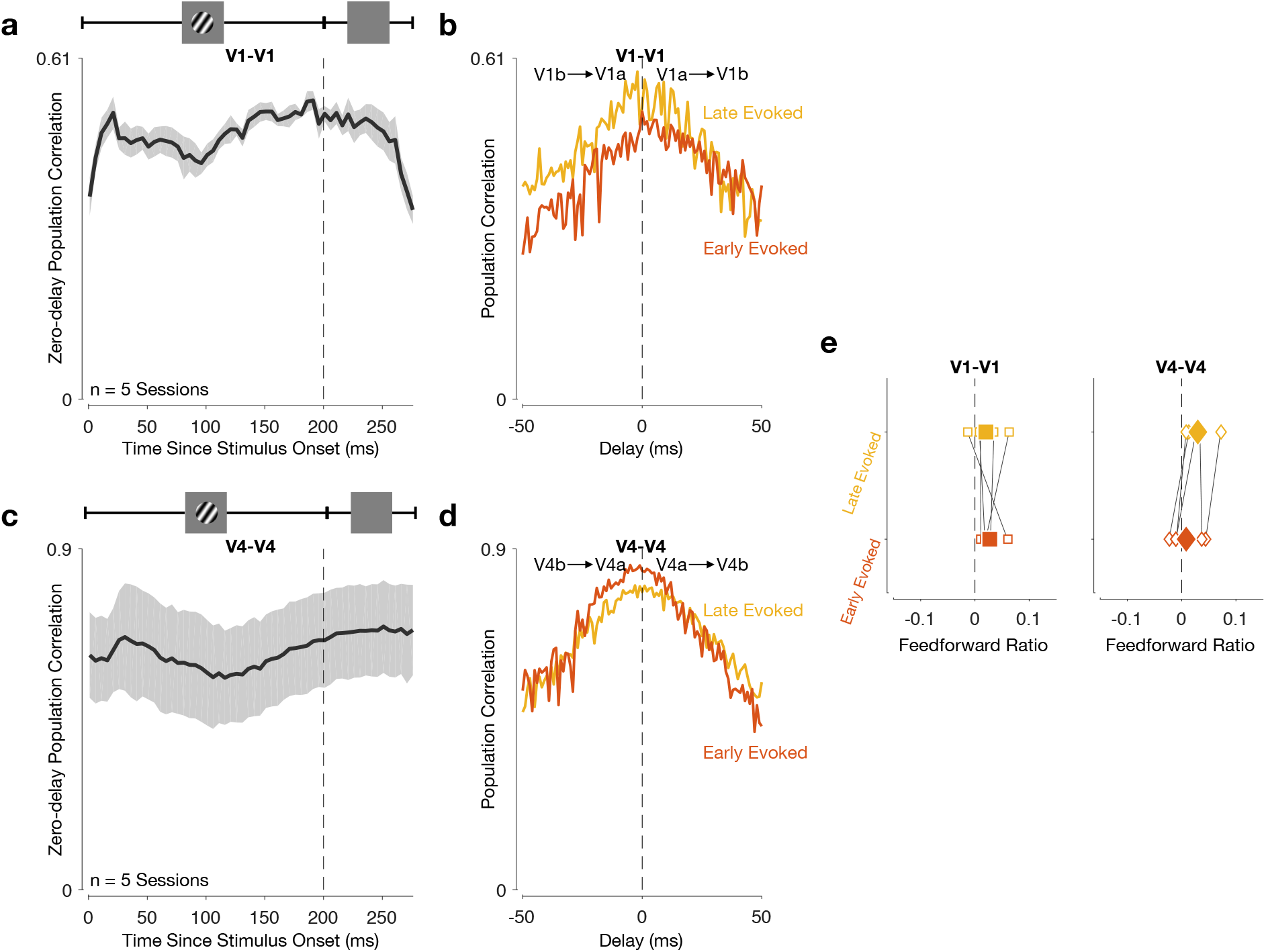
Interactions between subpopulations within V1 or V4 recorded in awake animals were neither feedforward-nor feedback-dominated. To test whether the effects in Fig. 4 were specific to inter-areal interactions, we randomly divided the neurons in each area into two subpopulations and computed population correlation functions between the subpopulations for each area. **(a)** V1-V1 zero-delay population correlation is constant throughout the trial. Shading indicates S.E.M. **(b)** V1-V1 population correlation functions for an example session (taken as the average across 25 random divisions into two V1 subpopulations). Same conventions as in Fig. 4b. **(c)** V4-V4 zero-delay population correlation is constant throughout the trial. Solid line shows average across all recording sessions. Shading indicates S.E.M. **(d)** V4-V4 population correlation functions for an example session (taken as the average across 25 random divisions into two V4 subpopulations). Same conventions as in Fig. 4b. **(e)** There are two key features of inter-areal interactions revealed in Fig. 4 which are not present for within-area interactions. First, the feedforward-dominated interaction shortly after stimulus onset (Fig. 4b,e) is absent here, and the correlation functions are centered at 0 ms delay (see panels b and d). Second, the transition from feedforward-to feedback-dominated interactions in the late evoked period (Fig. 4d) is also absent (average late evoked feedforward ratio, computed in the -50 to 50 ms delay range: 0.027 ± 0.008 SEM for V1-V1; 0.031 ± 0.011 SEM for V4-V4; one-sided paired Wilcoxon signed-rank test for difference between early evoked and late evoked activity across all 5 recording sessions, *p* = 0.41 for V1-V1; *p* = 0.97 for V4-V4).

**Supplementary Figure 5.**
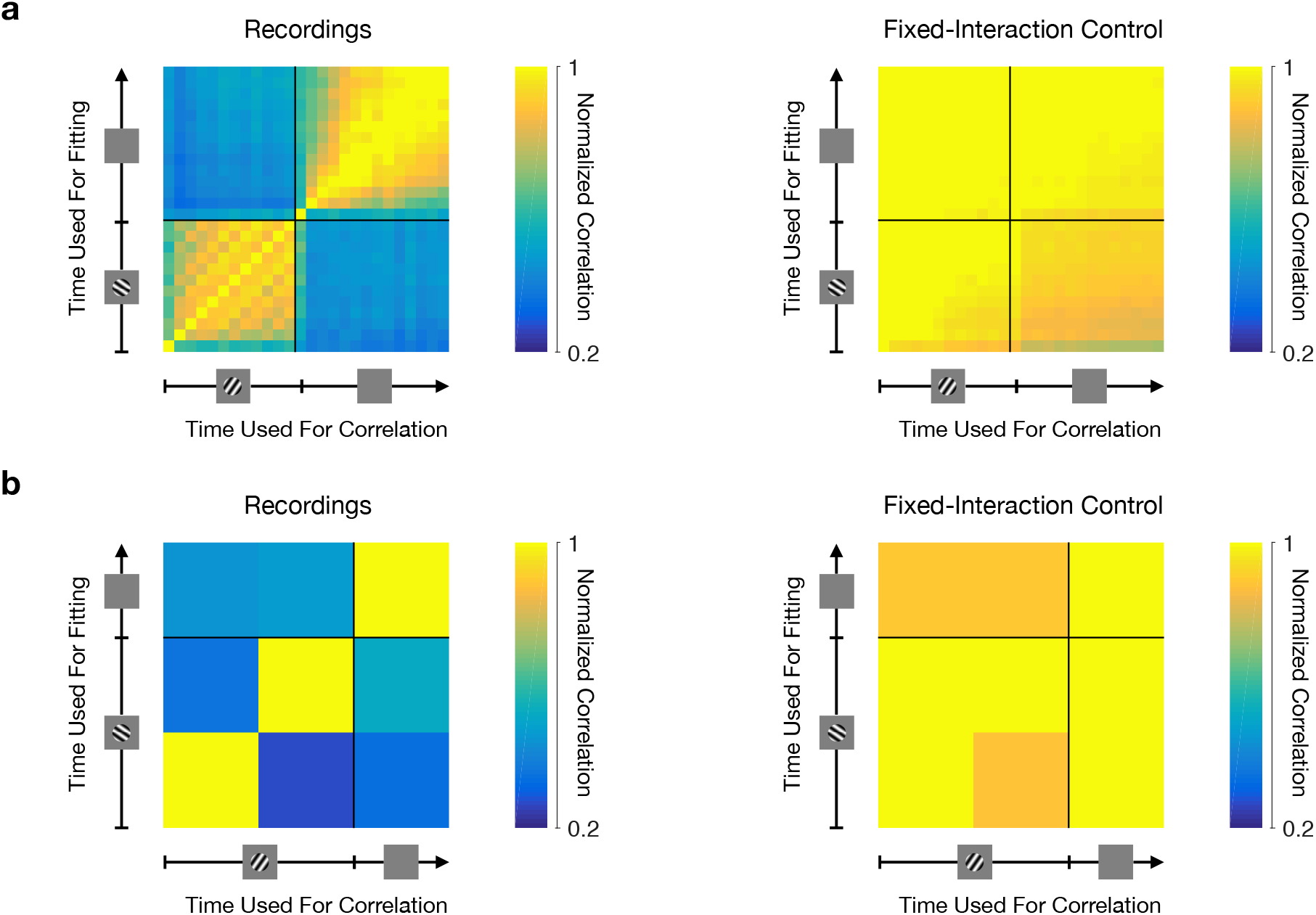
Ensuring that changes in activity patterns most related across areas cannot be ascribed to changes in the within-area population covariance structure. **(a)** We generated V1-V2 surrogate data that had approximately the same within-area covariance structure as the recorded data for each epoch, but for which the inter-areal interaction structure was held fixed (see Methods and Supplementary Information). For this synthetic data, our analysis identified a stable interaction structure (right, compare to left reproduced from Fig. 6d which is based on recorded activity). Same conventions as in Fig. 6d. **(b)** The same was true for the V1-V4 interactions (right, compare to left reproduced from Fig. 6e which is based on recorded activity). Same conventions as in Fig. 6e.

**Supplementary Figure 6.**
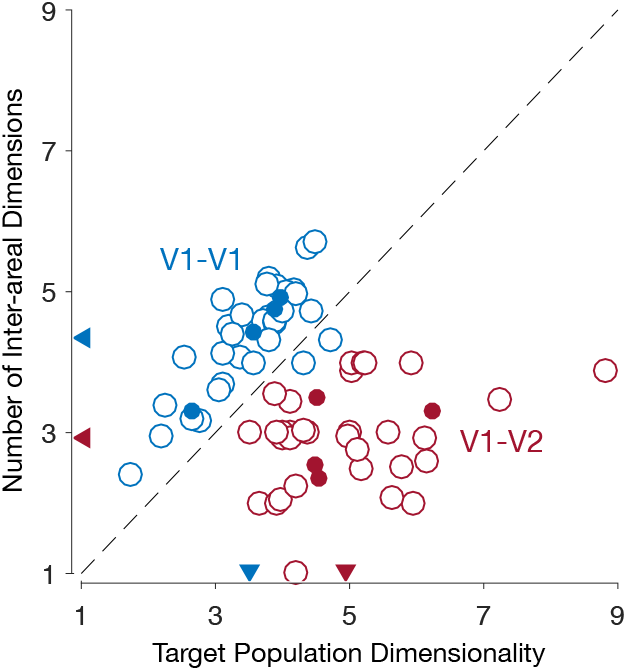
A communication subspace is evident when using Canonical Correlation Analysis (CCA) to characterize inter-areal interactions. We previously reported that the interaction between V1 and V2 was low dimensional (termed a communication subspace) using Reduced-Rank Regression (RRR)^29^. RRR is closely related to Canonical Correlation Analysis (CCA), which we employed in this work (for a review, see ref. 53, in press). One might wonder whether CCA also identifies a communication subspace between V1 and V2. We repeated the analysis in our previous work, using the same data that was analyzed there^29^, but using CCA instead of RRR to relate the activity across areas. To determine the number of dimensions involved in inter-areal interactions, we first evaluated the cross-validated log-likelihood curve for a probabilistic CCA model (pCCA), and picked the number of canonical dimensions that yielded the highest data likelihood. We then fit a pCCA model with the corresponding dimensionality using all trials, and computed the associated inter-areal covariance matrix. Finally, we used Singular Value Decomposition (SVD) to determine the smallest number of dimensions that captured 95% of the inter-areal covariance, and used that number as our estimate of the number inter-areal dimensions.

As in our previous work^29^, we found that fewer dimensions were required to characterize inter-areal interactions (V1-V2; red triangle on vertical axis) than within-area interactions (V1-V1; blue triangle on vertical axis). In contrast to Fig. 3, where we identified a single significant canonical pair for each epoch and time delay, here we identify on average close to 3. This is largely due to the larger binning windows used here (100 ms vs. 1 ms in Fig. 3). Importantly, the lower number of dimensions required to account for inter-areal interactions, compared to within-area interactions, was not a result of lower dimensional activity in the V2 population, as the population activity dimensionality was higher in V2 than in the held-out V1 populations (compare blue and red triangles on horizontal axis). Moreover, the number of predictive dimensions identified by RRR was highly correlated with the number of canonical dimensions identified by CCA (Pearson correlation coefficient *r*^2^ = 0.89 across all datasets; not shown). Open circles corresponds to each dataset, filled circles denote mean across datasets for each recording session. Triangles denote mean across all recording sessions.

## Supplementary Information

### Characterizing changes in the interaction structure

What constitutes a change in the interaction structure? In other words, how can we evaluate whether or not different activity patterns are involved in inter-areal interactions during different trial epochs? Using Canonical Correlation Analysis (CCA, see Methods) to characterize inter-areal interactions, one might wonder whether changes in the canonical dimensions across two epochs are a good indication of a change in the interaction structure. Here, we show that directly levering the canonical dimensions to test for changes in the interaction structure can be misleading, and propose an alternative approach based on the probabilistic CCA (pCCA) model^64^.

Suppose we identify *q* pairs of canonical dimensions, and represent them as the columns of matrices A_*q*_ and B_*q*_, which have dimensions *p*_*x*_ × *q* and *p*_*y*_ × *q*, respectively, where *p*_*x*_ and *p*_*y*_ are the number of recorded neurons in each of the two areas. The column space of each matrix defines a subspace in each area within which activity is most correlated across areas. If one seeks to compare the canonical dimensions identified during two trial epochs, one possibility is to compare the column spaces of matrices A_*q*_ and B_*q*_ for two different epochs.

There is, however, a potential problem with using this approach to ask whether there was a meaningful change in the inter-areal interaction structure. We can illustrate this issue by considering data generated from a pCCA model. pCCA is defined by the following equations:

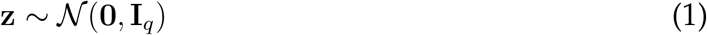

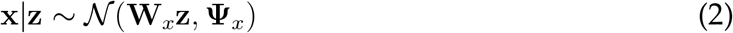

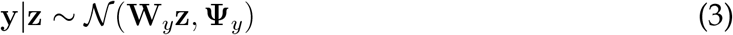

where z is a *q* × 1 latent variable, x and y correspond to the neuronal activity recorded in each of two cortical areas, with dimensionalities *p*_*x*_ × 1 and *p*_*y*_ × 1, respectively, *p*_*x*_ and *p*_*y*_ are the number of neurons recorded in each area, and *q*≤ min(*p*_*x*_, *p*_*y*_). The identity matrix **I**_*q*_ has dimensions *q* × *q*. The mapping matrices **W**_*x*_ and **W**_*y*_ have dimensions *p*_*x*_ × *q* and *p*_*y*_ × *q*, respectively. The covariance matrices **Ψ**_*x*_ and **Ψ**_*y*_ have dimensions *p*_*x*_ × *p*_*x*_ and *p*_*y*_ × *p*_*y*_, respectively. We assume, without loss of generality, that x and y are mean-centered. CCA and pCCA return the same correlation values, so both methods result in the same population correlation functions. The advantage of pCCA here is that it provides us with a more complete description of the fitted model and its underlying assumptions.

According to pCCA’s graphical model (Fig. S1a), we can describe the observed activity in each area as having an “across-area” component and a “within-area” component. The across-area component emerges via the linear mapping between the shared latent variable z and each observed variable, x and y. This mapping is defined by the matrices **W**_*x*_ and **W**_*y*_. The within-area components are defined to be Gaussian with unconstrained covariance matrices**Ψ** _*x*_ and**Ψ** _*y*_.

The relationship between the column spaces of the matrices A_*q*_ and B_*q*_ computed by classical CCA and the parameters of the pCCA model is given by:

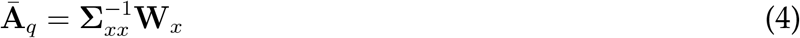

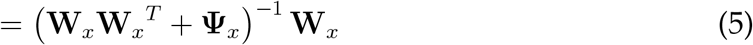

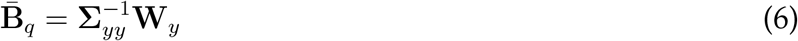

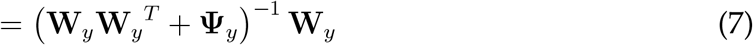

where Ā _*q*_ and 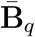 have the same column space as A_*q*_ and B_*q*_, respectively (see the “Relationship between CCA and pCCA” section below). This shows that the subspaces spanned by the canonical dimensions in each area depend on the within-area noise parameters **Ψ**_*x*_ and **Ψ**_*y*_. Thus, changes to the within-area components lead to changes in the subspaces spanned by the canonical dimensions, even if the across-area components remain fixed. Measuring changes in the interaction structure by measuring to what extent the subspaces spanned by the canonical dimensions differ would thus lead us to conclude that across-area interaction structure had changed, even though only the within-area components were altered.

We can gain further intuition into the pCCA model by inspecting the joint covariance matrix (Fig. S1b) The covariance for each area, Σ_*xx*_ (Σ_*yy*_), is composed of an across-area component, **W**_*x*_**W**_*x*_^*T*^ (resp. W_*y*_W_*y*_^*T*^) and within-area component,**Ψ** _*x*_ (resp. **Ψ**_*y*_). Figure S1c illustrates this covariance decomposition for one of the areas (ellipses represent each covariance component). For the across area covariance, however, we have 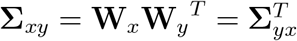 Thus, the across-area covariance structure is solely determined by the linear mapping matrices **W**_*x*_ and **W**_*y*_.

Given that the across-area component in the pCCA model is solely determined by the mapping matrices **W**_*x*_ and **W**_*y*_, we can quantify changes in the interaction structure by comparing those matrices for different trial epochs. We will take this approach, and use a pCCA model to estimate the **W**_*x*_ and **W**_*y*_ matrices, and in turn use changes in these matrices to detect changes in the interaction structure. We need to first define how to measure differences between the W_*x*_ and **W**_*y*_ matrices estimated at different times during the trial. As mentioned above, **W**_*x*_ and **W**_*y*_ are underdetermined, so an element by element comparison (e.g., the Frobenius norm of the difference between two **W**_*x*_ matrices fit at different epochs in the trial) is not suitable. We defined our difference metric to be based on differences between the column spaces of **W**_*x*_ and **W**_*y*_, i.e., our measure of how much the interaction structure changes across different epochs is only sensitive to changes in the subspaces spanned by the dimensions along which activity is related across areas. To be conservative, we will not consider scaling and affine transformations of these dimensions (which do not change the subspace spanned by these dimensions) as changes to the interaction structure, although they might reflect interesting changes for other analysis goals.

Specifically, we will measure differences between the column spaces of **W**_*x*_ and **W**_*y*_ by comparing the inter-area correlation these subspaces account for. To compare the interaction structure identified during two epochs in the trial, indexed by *t* and *t*′, we first fit a pCCA model at each epoch, yielding parameters 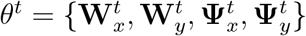 and 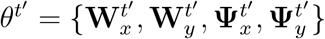 We then ask: given the observed (sample) within-area covariance matrices at time *t*′, 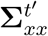 and 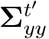, how correlated would the activity across areas be if instead of the estimated matrices 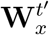 and 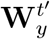, the interaction was instead described by matrices 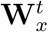 and 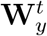? More specifically, how much do across-area correlations change if we replace the column space of 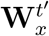 and 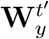 by the column space of 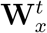 and 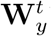? We can use equations 4 and 6 to compute the subspace spanned by the canonical dimensions induced by 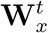 and 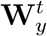 (see the “Relationship between CCA and pCCA” section below):

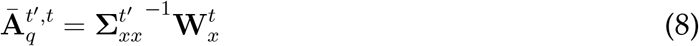

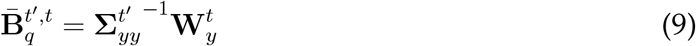

and measure the amount of across-area correlation captured by 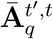 and 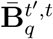 We then compare that amount of across-area correlation to the correlation that would have resulted from using 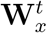 and 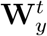 (i.e., the across-area correlation captured by 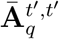 and 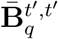). The results of this analysis are shown in Fig. 6. Note that both 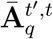 and 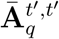 (resp. 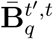 and 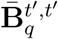) are computed using the same covariance matrix 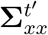(resp. 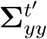. Thus, any differences between 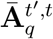 and 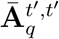 (resp.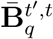 and 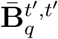) are the result of differences between 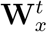 and 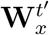 (resp.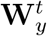 and 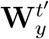). Specifically, differences between the column spaces of 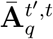 and 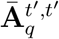 (resp.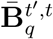 and 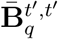 are the result of differences between the column spaces if 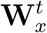 and 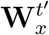 (resp 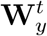 and 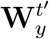; see “Relationship between CCA and pCCA” section below). Algorithm (1) describes this process in detail.

**Algorithm 1:** Comparing inter-area interaction structure across time

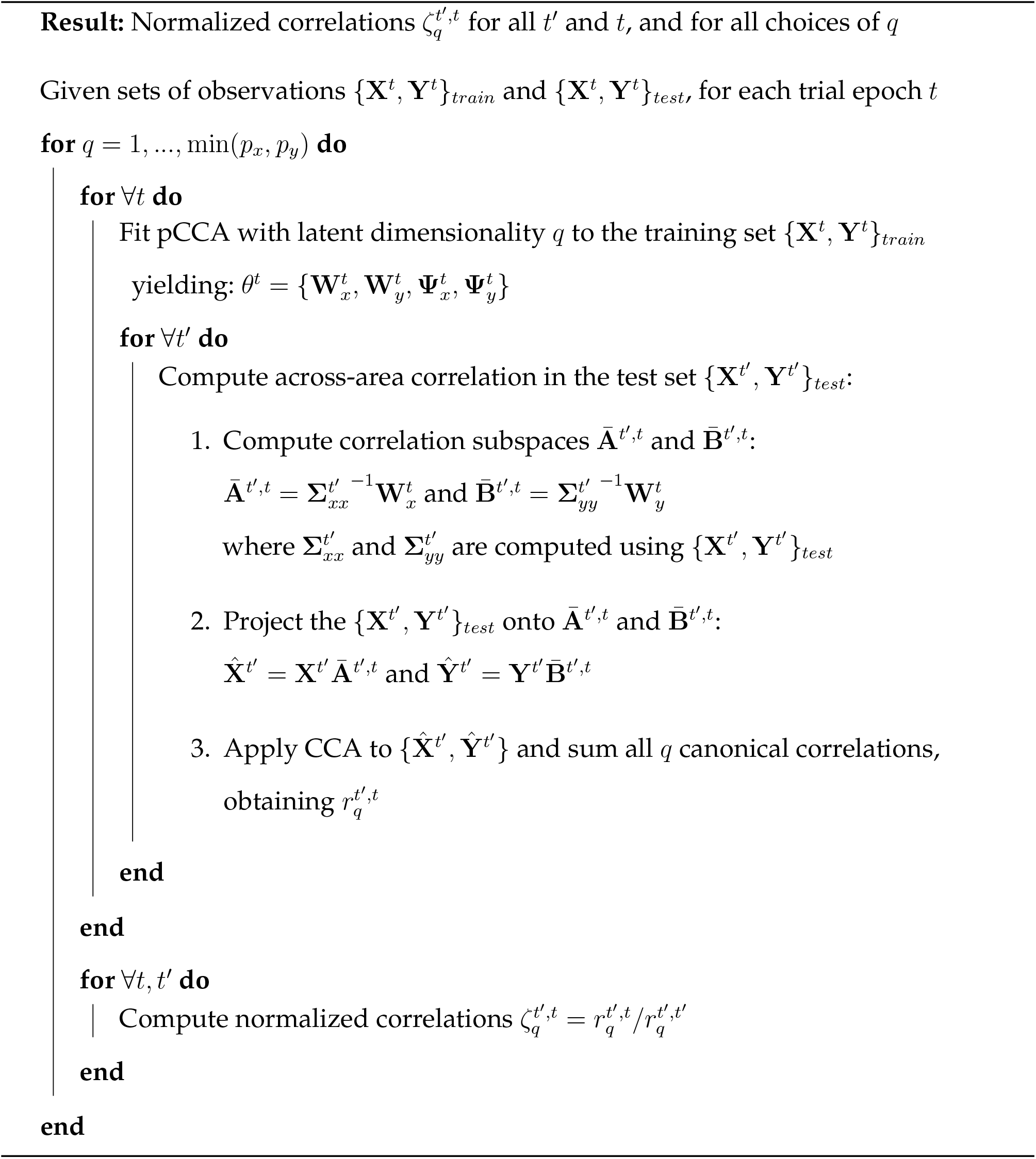

Algorithm (1) describes a simple train/test split, but it is easy to generalize this procedure to run within a k-fold cross-validation scheme (and then average the normalized correlations across test folds). In the current study, we employed 10-fold cross-validation.

### Fixed interaction structure control

The analysis described in Algorithm (1) was designed to be sensitive only to the column spaces of the **W**_*x*_ and **W**_*y*_ matrices. To empirically test that our analysis is insensitive to changes in the remaining pCCA model parameters (i.e., that the changes reported in Fig. 6 are solely due to changes to **W**_*x*_ and **W**_*y*_), we devised a control based on the following intuition: if we analyze data where the across-area component is held fixed while the within-area component changes, our method (if it works as we expect it to) should indicate that there is no change in interaction between areas. In other words, if we keep the column spaces constant across epochs, we should find that all normalized correlations will be close to 1 (i.e., we identify the same column spaces throughout the trial). To carry out this control analysis, we generated surrogate data that was as similar as possible to the observed activity (in terms of the first and second order statistics, number of trials and number of observed neurons), but with fixed column spaces for the mapping matrices W_*x*_ and **W**_*y*_.

To achieve this, we first fit a pCCA model to the recorded neural activity, across all epochs, obtaining matrices **W**_*x*_ and **W**_*y*_. We then choose matrices 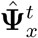 and 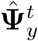 for each epoch such that 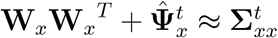 and 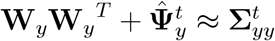 for each epoch *t*. Note that **W**_*x*_ and **W**_*y*_ are the same for all epochs. Figure S2 illustrates this for two epochs *t* and *t*′.

For 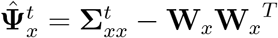 to be a proper covariance matrix, it must be positive definite, which is not guaranteed to be the case (similarly for 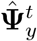). A simple way to ensure that is positive definite is to scale **W**_*x*_ and **W**_*y*_ appropriately for each time step, as this operation does not change their column spaces. Algorithm (2) describes the surrogate data generation process in detail.

We found that fixing the column spaces of the **W**_*x*_ and **W**_*y*_ in this way led pCCA to identify fixed columns spaces across all epochs (Supplementary Fig. 5a,b), indicating the results in Fig. 6 are not driven by changes in the within-area components but rather by changes in the inter-areal interaction structure.

**Algorithm 2:** Creating surrogate data with a fixed interaction structure

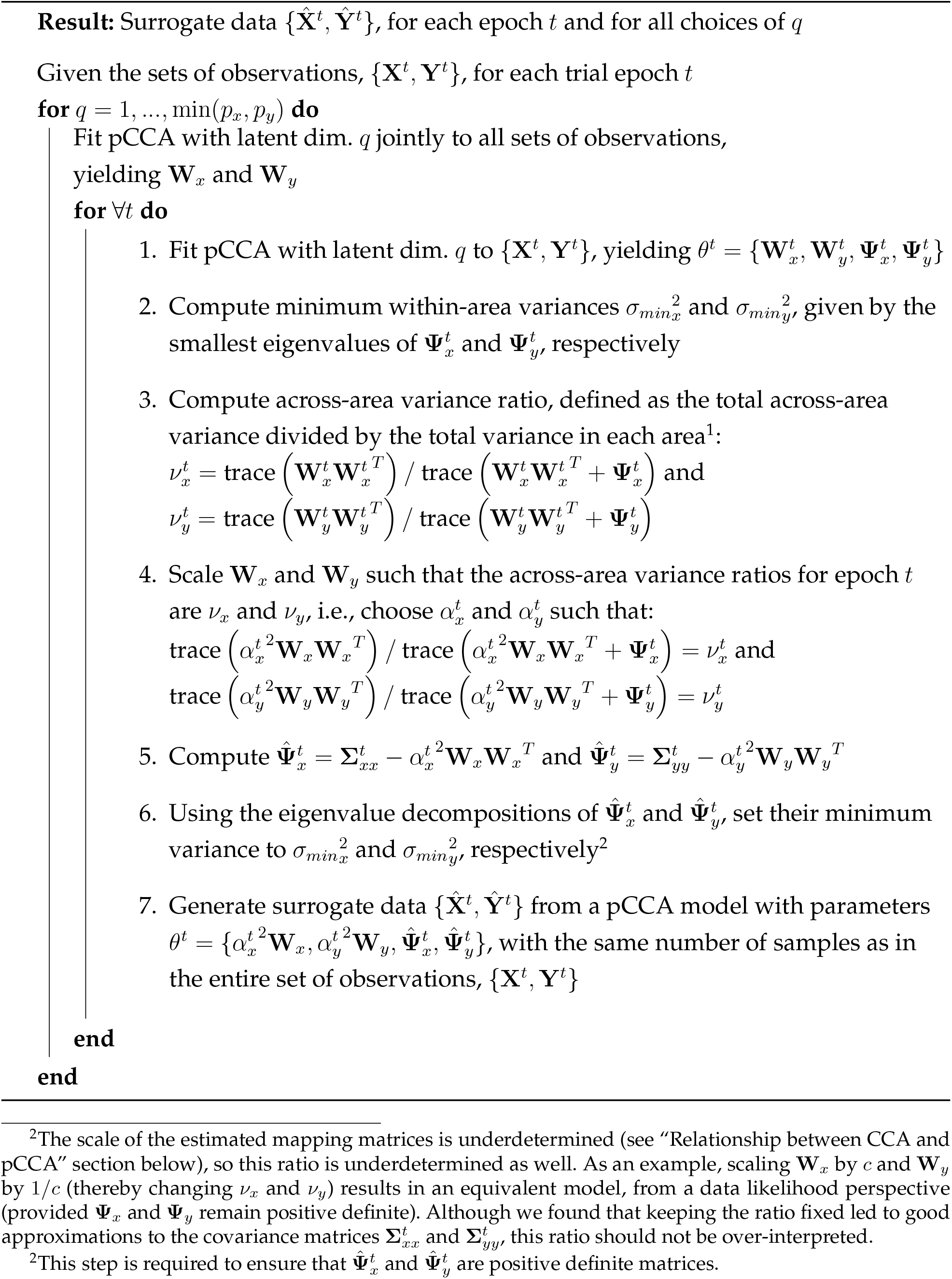

### Relationship between CCA and pCCA

The correspondence between the classical formulation and the probabilistic variant of CCA was developed by Bach and Jordan^64^. Specifically, they showed that any maximum likelihood solution derived using the probabilistic model (equations 1-3) corresponds to the same set of canonical dimensions identified using classical CCA. In other words, the data likelihood function for pCCA has infinitely many global optima, where all local optima are also global optima, and all global optima correspond to the same set of canonical dimensions. In particular, if we define the top *q* canonical dimensions identified for each area by classical CCA as **A**_*q*_ (a *p*_*x*_ × *q* matrix) and **B**_*q*_ (a *p*_*y*_ × *q* matrix), the relationship between the canonical dimensions and the linear mapping matrices from pCCA is given by:

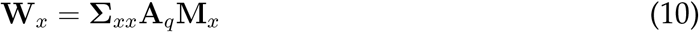

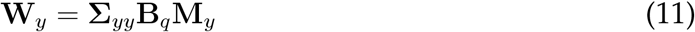

where **M**_*x*_ and **M**_*y*_ are arbitrary *q* × *q* matrices such that **M**_*x*_**M**_*y*_^*T*^ = **P**_*q*_ and the spectral norms of **M**_*x*_ and **M**_*y*_ are smaller than one. **P**_*q*_ is a diagonal matrix containing the first *q* canonical correlations. As an example, 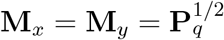 satisfies these constraints. Any suitable choice of **M**_*x*_ and **M**_*y*_ corresponds to a global maximum of the data likelihood. The link between CCA and pCCA is similar to that between PCA and pPCA, and the derivation of this connection largely follows that originally developed for PCA and pPCA^65,66^.

In particular, the fact that **M**_*x*_ and **M**_*y*_ are underdetermined means that **W**_*x*_ and **W**_*y*_ are not uniquely defined when fitting pCCA, i.e., there are many choices of **W**_*x*_ and **W**_*y*_ that result in the same canonical dimensions, and maximizing the data likelihood can return any such choices. Importantly, these **W**_*x*_ (**W**_*y*_) matrices all have the same column space, i.e., multiplication by **M**_*x*_ (resp. **M**_*y*_) does not change the column space of **W**_*x*_ (resp. **W**_*y*_).

Given two matrices **W**_*x*_ and **W**_*y*_ found by maximizing the data likelihood, we cannot directly compute **A**_*q*_ and **B**_*q*_ from these matrices alone, since we don’t know which **M**_*x*_ and **M**_*y*_ the particular solution we found corresponds to. However, since all the consistent **W**_*x*_ (**W**_*y*_) matrices have the same column space, we can find the column spaces of **A**_*q*_ and **B**_*q*_ by computing matrices Ā _*q*_ and 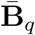 (equations 4 and 6), since the column space of Ā _*q*_ 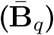 is the same as the column space of **A**_*q*_ (resp. **B**_*q*_) (see Lemma 1 below). Note that the column space of **A**_*q*_ (resp. **B**_*q*_) is the subspace of x (y) spanned by the canonical dimensions found by classical CCA. The relationship above indicates that the subspace spanned by the canonical dimensions in **A**_*q*_ (**B**_*q*_) depends on the column space of **W**_*x*_ (resp. **W**_*y*_) and on Σ_*xx*_ (resp. Σ_*yy*_). In particular, if Σ_*xx*_ (Σ_*yy*_) is held fixed, the column space of **A**_*q*_ (resp. **B**_*q*_) is solely determined by the column space of **W**_*x*_ (resp. **W**_*y*_; see Lemma 2 below). This observation forms the basis for Algorithm 1, where we ask how well a pCCA model fit to epoch *t* (yielding 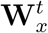 and 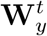) captures correlations at epoch *t*′.

#### Lemma 1

**Ā** _*q*_ 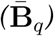 *and* **A**_*q*_ *(resp*. **B**_*q*_*) have the same column space*.

*Proof*. We will show that **Ā** _*q*_ and **A**_*q*_ have the same column space. The proof for 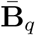 and **B**_*q*_ is identical. Starting with equation 10:

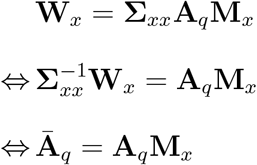

where we used the fact that Σ_*xx*_ is a square positive definite matrix. So long as M_*x*_ is a full rank matrix (i.e., the first *q* canonical correlations are non-zero), Ā _*q*_ = A_*q*_M_*x*_ and A_*q*_ have the same column space.

#### Lemma 2.

*If* Σ_*xx*_ (Σ_*yy*_*) is held fixed, the column space of* **A**_*q*_ *(resp*. **B**_*q*_*) is solely determined by the column space of* **W**_*x*_ *(resp*. **W**_*y*_).

*Proof*. We will show that the column space of **A**_*q*_ depends solely on the column space of **W**_*x*_ if Σ_*xx*_ is held fixed. The proof for B_*q*_ is identical. Using the compact singular value decomposition **W**_*x*_ = **UDV**^*T*^, and inserting it into equation 10:

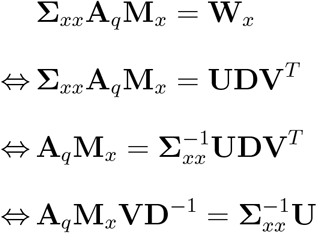

where we used the fact that Σ_*xx*_ is a square positive definite matrix. As long as **M**_*x*_ is a full rank matrix (i.e., the first *q* canonical correlations are non-zero), **M**_*x*_**VD**^-1^ is a square full rank matrix, and thus **A**_*q*_**M**_*x*_**VD**^-1^ and **A**_*q*_ have the same column space. So as long as Σ_*xx*_ is held fixed, the column space of **A**_*q*_ only depends on U, which is a basis for the column space of **W**_*x*_. In other words, if we change **W**_*x*_, only the changes to U (its column space), and not changes to D or V, affect the column space of A_*q*_.

**Figure S1.**
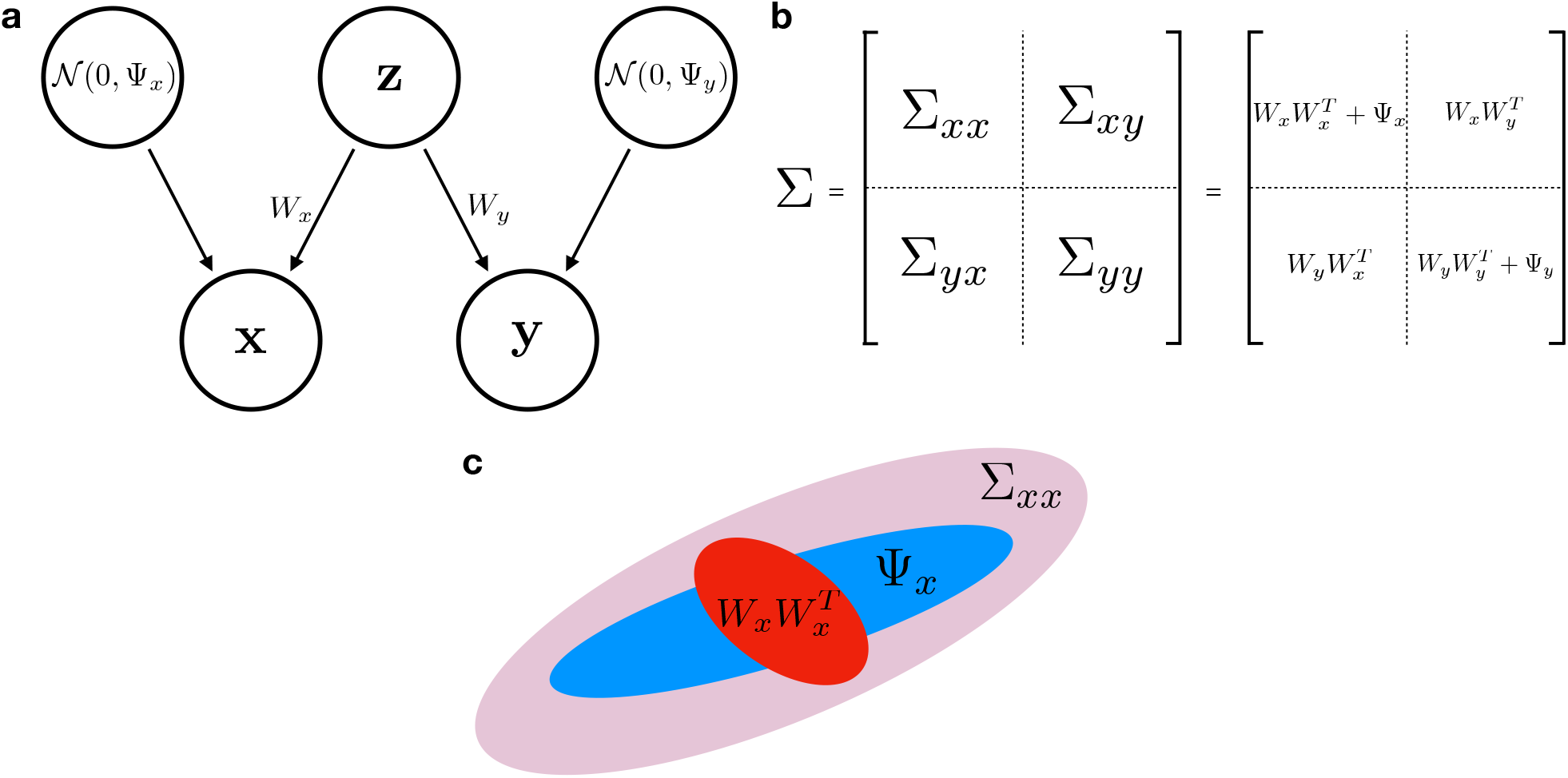
Probabilistic canonical correlation analysis (pCCA) **(a)** pCCA’s probabilistic graphical model. **(b)** Summary of the relationship between the data covariance matrices and the pCCA model parameters. **(c)** Graphical representation of the covariance decomposition under a pCCA model for one of the two populations. Red ellipse represents the across-area component; blue ellipse represents the within-area component; pink ellipse represents the total covariance in this area.

**Figure S2.**
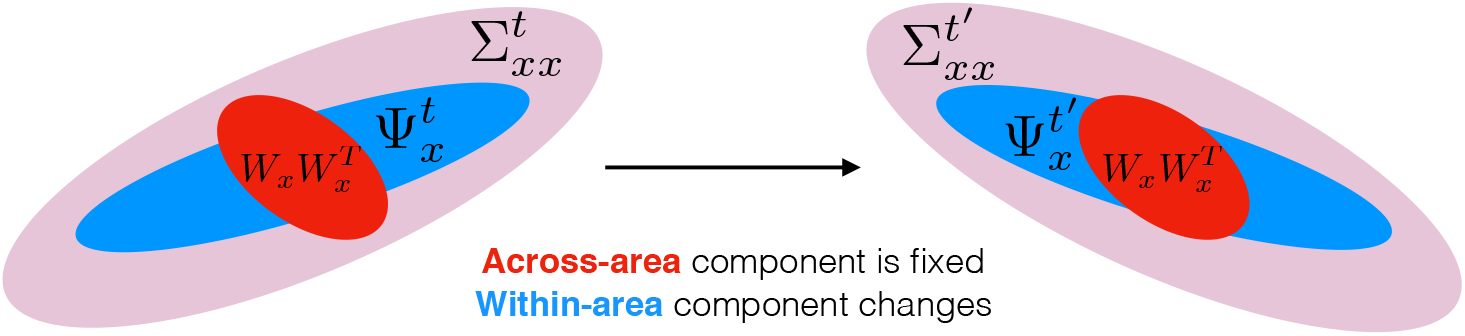
Changing the total covariance in one of the areas while keeping the across-area component fixed. Same conventions as in Fig. S1c.

